# Skin-resident immune cells actively coordinate their distribution with epidermal cells during homeostasis

**DOI:** 10.1101/2021.01.08.425932

**Authors:** Sangbum Park, Catherine Matte-Martone, David G. Gonzalez, Elizabeth A. Lathrop, Dennis P. May, Cristiana M. Pineda, Jessica L. Moore, Jonathan D. Boucher, Edward D. Marsh, Axel Schmitter, Katie Cockburn, Yohanns Bellaïche, Valentina Greco

## Abstract

Our organs consist of multiple cell types that ensure proper architecture and function. How different cell types coexist and interact to maintain their homeostasis *in vivo* remain elusive. The skin epidermis comprises mostly epithelial cells, but also harbors Langerhans cells (LCs) and Dendritic Epidermal T cells (DETCs). In response to injury or infection, LCs and DETCs become activated and play critical immunological roles. During homeostasis, they coexist with epithelial cells in the basal layer of the epidermis. Whether, and how, distributions of LCs and DETCs are regulated during homeostasis is unclear. Here, we addressed this question by tracking LCs, DETCs and epithelial basal cells over time within the skin of live adult mice. We show that LCs and DETCs maintain their overall position despite continuous turnover of neighboring basal epithelial stem cells. Moreover, LCs and DETCs rapidly and maximally explore basal epithelial cell junctions through their dendritic extensions. Altering the epithelial cell density triggers corresponding changes in the immune cell density, but not vice versa, suggesting that epithelial cells determine immune tissue composition in the epidermis. Moreover, LCs and DETCs are organized in a tiling pattern that is actively maintained. When LCs or DETCs are ectopically removed, neighboring epidermal LCs or DETCs, respectively, move into the emptied spaces and re-establish the tiling pattern. Finally, LCs require the GTPase Rac1 to maintain their positional stability, density and tiling pattern. Overall, we discovered that epidermal cells regulate the density of immune cells during homeostasis, and that immune cells actively maintain a non-random spatial distribution, reminiscent of neuronal self-avoidance. We propose that these cellular mechanisms provide the epidermis with an optimal response to environmental insults.

## Main text

The skin epidermis is the outermost layer of our body and acts as a barrier to protect us from the environment. The epidermis is mainly composed of epithelial cells, which are continuously replenished by epithelial stem cells that reside in the basal layer [1–6]. These basal epithelial cells are closely intermingled with two main populations of skin-resident immune cells, Langerhans cells (LCs) and dendritic epidermal T cells (DETCs) [7–9]. Many studies have advanced our understanding of the immunological functions of skin-resident immune cells during injury or inflammation, as well as of the signaling supporting their surveillance role in the epidermis [10–27]. Additionally, previous work has also captured immune cells actively surveilling the surface of the skin epidermis [28–34]. Nevertheless, it is not known how these skin-resident immune cells regulate their homeostasis within the continuously regenerating epidermis, and to what extent the dynamic behaviors of these diverse cell types are coordinated.

To investigate the behaviors of LCs and DETCs as they coexist with epithelial cells in live mice, we developed a mouse line that labels the three main cell types within the skin epidermis using distinct fluorescent markers (**Figure 1a**). These mice contain a huLangerin-CreER;tdTomato reporter [35, 36] that labels LCs with tdTomato after tamoxifen injection. In addition, they express GFP under the control of the CX3CR1 promoter [37], which labels DETCs in the epidermis, as well as Keratin14-Histone2B Cerulean [38, 39] which labels all epithelial stem cell nuclei and derived lineages. We utilized our previously established intravital imaging approach [40, 41] to visualize all three populations (LCs, DETCs and epithelial cells). By acquiring stacks along the whole apical-basal axis of the skin, we uncovered that LCs and DETCs are embedded within the basal stem cell layer of the epidermis (**Figure 1a-c, Supplementary Video 1**). The dendritic radial morphology of these immune cells prompted us to ask how many epithelial basal cells they contact. To this end we quantified the immune/epithelial basal cell contacts in different geographic skin regions: the ear, which harbors LCs and DETCs, and the paw, which harbors only LCs. In both regions, nearly all epithelial basal cells were in contact with either LCs and/or DETCs. This reveals that LCs and DETCs extensively cover the basal stem cell layer (**Figure 1d-e**).

**Figure 1.**
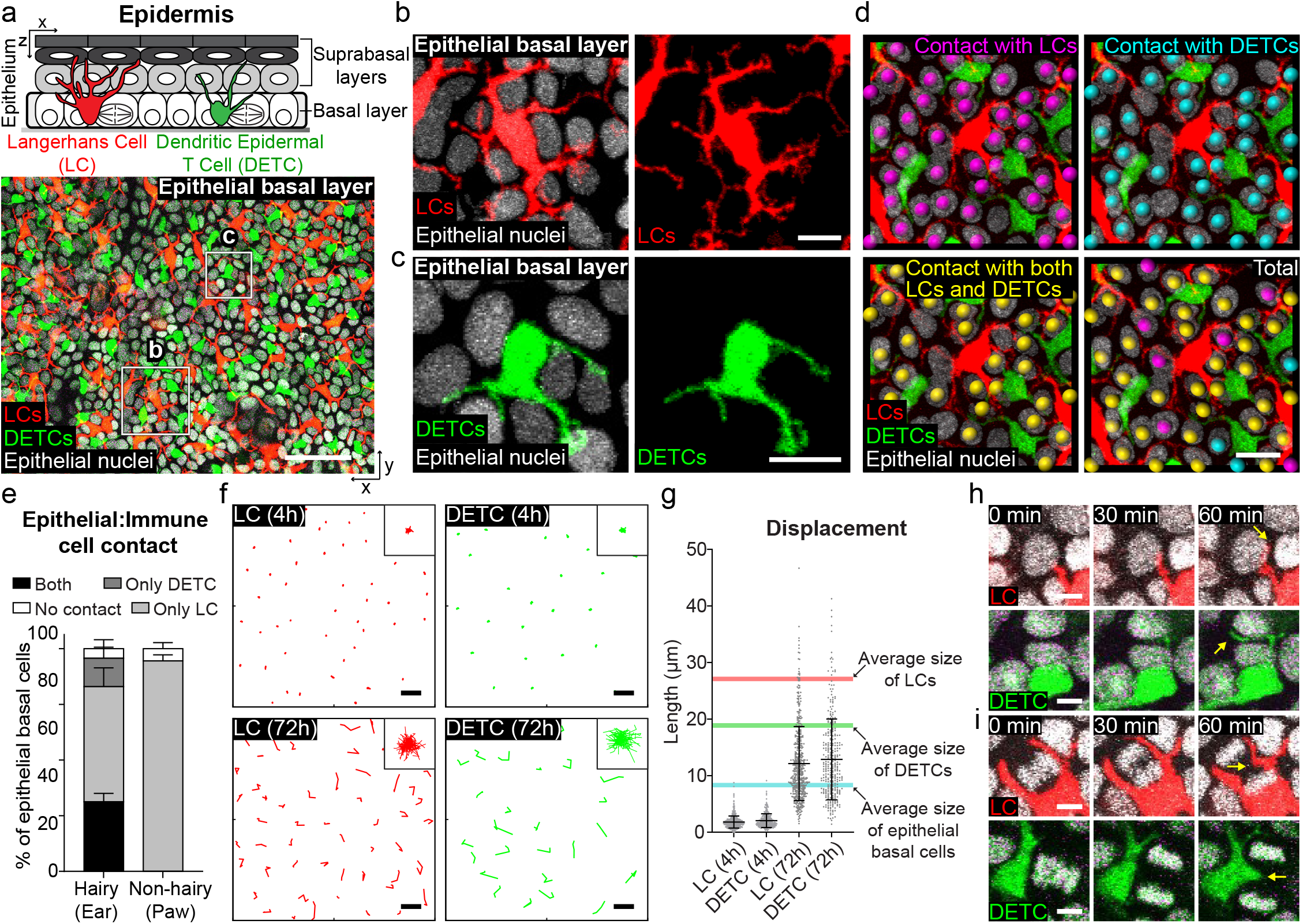
LCs and DETCs maximally cover and adapt to changes of neighboring epithelial basal cells during homeostasis. (a) Langerhans cells (LCs) and dendritic epidermal T cells (DETCs) coexist with epithelial basal cells and differentiated cells in the epidermis. The illustration shows an x-z view of the whole mouse epidermis with the basal layer highlighted in gray (top panel). The bottom image is an x-y view of the basal layer, which contains LCs, DETCs, and exclusively epithelial basal cells (bottom panel) (red, LCs; green, DETCs; white, nuclei of epithelial cells; representative images from 3 *huLangerin-CreER; Rosa-stop-tdTomato; CX3CR1-GFP; Kl4-H2BCerulean* mice). Scale bar, 50 μm. (b, c) LCs and DETCs are embedded between epithelial basal cell neighbors via their dendritic radial morphology (red, LCs; green, DETCs; white, nuclei of epithelial cells; representative images from 3 mice). Scale bar, 10 μm. (d) As the epithelial basal cell nucleus/cytoplasmic ratio is high, cell-nucleus contact between immune and epithelial basal cells is used for an approximation of cell-cell contact. The direct contact of epithelial basal cells with LCs is visualized with magenta balls, DETCs with cyan balls and/or both LCs and DETCs with yellow balls (representative images from 3 mice). Scale bar, 20 μm. (e) Quantification of cell-cell contact between basal epithelial cell nuclei and LCs only (light gray), DETCs only (dark gray), both LCs and DETCs (black) or neither (white) (n=3 mice respectively). (f) Displacement track analysis of individual LCs (in red) and DETCs (in green; upper panel, 4 hour timelapse; bottom panel, revisit every 24 hours for 3 days; representative images from 3 mice). Insets in upper right corner show the overlay of all the individual tracks (rose plots) which represent the collective displacement of all tracked LCs and DETCs. Scale bar, 20 μm. (g) Quantification of displacement length of displacement tracks (n=3 mice respectively). The red line is the average size of LCs (27.07 μm, 574 cells from 3 mice), the green line is the average size of DETCs (18.86 μm, 390 cells from 3 mice), and the blue line is the average size of epithelial basal cells (8.30 μm, 600 cells from 3 mice) (h, i) Images from a time-lapse show that the dendrites of LCs and DETCs continuously explore junctional spaces with neighboring epithelial basal cells including those newly formed after division (h, non-dividing cells; i, dividing cells; representative images from 3 mice). Scale bar, 5 μm.

Epithelial stem cells continuously remodel the basal layer, via cell gain due to cell division and cell loss via differentiation and delamination. Each cell undergoes one of these behaviors on average every 2.5 days [41]. To examine how LCs and DETCs behave within these dynamic basal epithelial neighborhoods, we followed the same immune cells over time by recording time-lapse movies for 4 hours and also by revisiting the same area every 24 hours over 3 days [40, 42, 43]. We analyzed their movement by creating displacement tracks and found that the cell bodies of both LCs and DETCs move slowly over 72 hours **(Figure 1f, g**). Time-lapse movies showed that, in contrast to their slowly moving cell bodies, the dendrites of both LCs and DETCs actively extend and retract between several of their basal epithelial cell neighbors and between newly formed junctional spaces, such as the ones between just born daughter cells upon cell divisions **(Figure 1h, i, Supplementary video 2)**. Together, these data suggest that LCs and DETCs overall maintain positional stability over time. They do so by slowly adjusting their cell body location in response to the remodeling of their basal epithelial cell neighborhood, while constantly exploring this changing stem cell environment via dynamic dendritic movements.

Given the extensive direct contact between LCs and DETCs and epithelial cells in the basal layer, we asked whether their densities were coordinated. To investigate this question, we first turned to different skin regions, ear and paw, which harbor distinct epithelial densities [41]. Quantifications across these regions showed that the density of LCs was increased in the paw compared to the ear, correlating to the increased density of epithelial basal cells in the paw **(Figure 2a-c)**. Interestingly, the ratio between LCs and epithelial basal cells remained constant across these different skin regions (**Figure 2d**).

**Figure 2.**
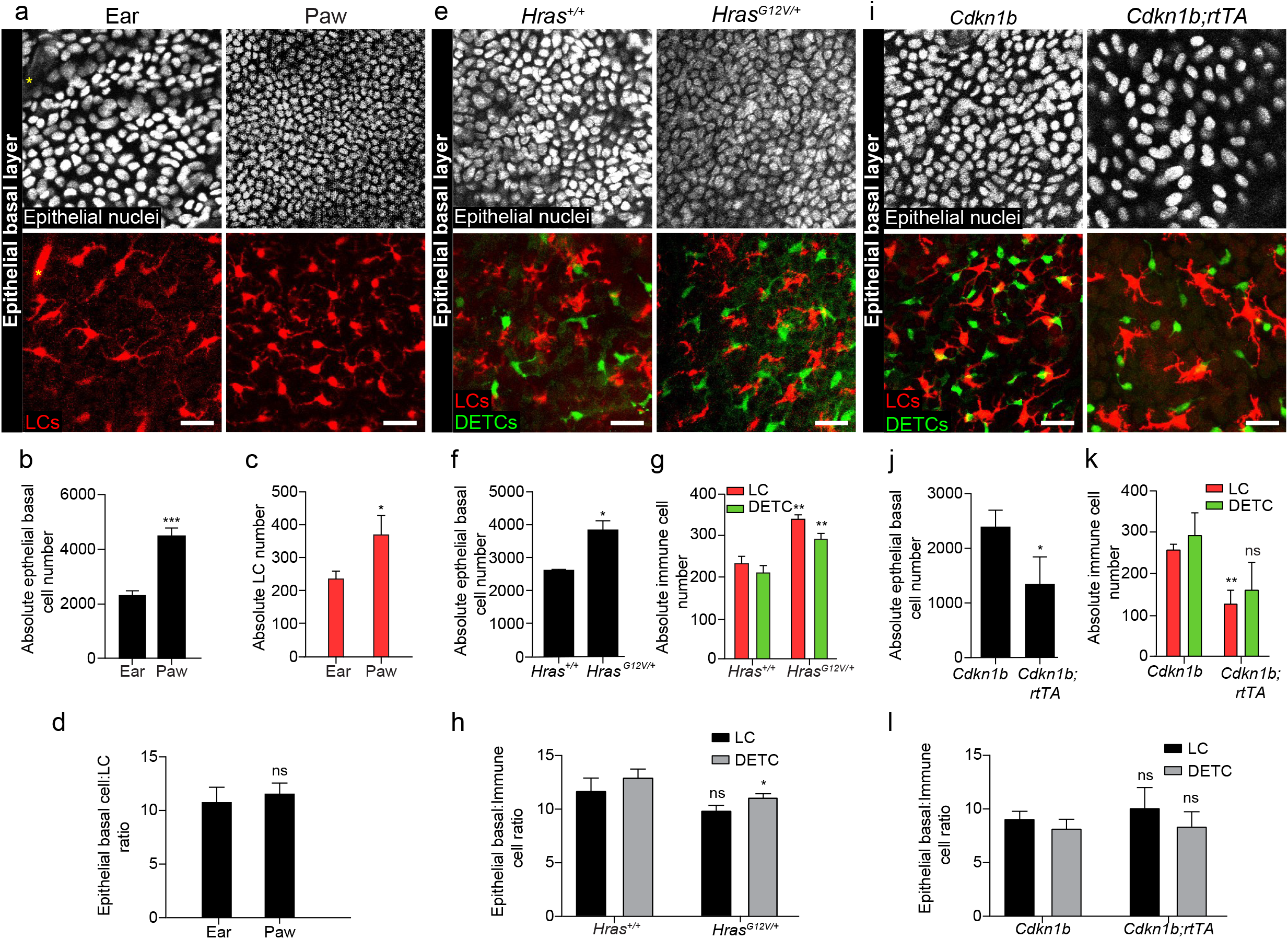
Immune cell density is regulated by epithelial basal cell density. (a) The paw has higher densities of epithelial basal as well as immune cells in the basal layer of epidermis (right) when compared to the ear (left) (representative images from 3 mice). DETCs are not present in the paw epidermis in the utilized models. Epithelial nuclei are white (top panel). LCs are red (bottom panel). Yellow asterisk denotes hair follicles. Scale bar, 30 μm. (b,c) Cell number quantifications were compared between ear and paw and found to be significant for both epithelial basal cells and LCs. Epithelial basal cell number (b) and LC number (c) in 0.25 mm^2^. * *p*<0.05 and *** *p*<0.001, unpaired Student’s t-test (n=3 mice respectively). (d) The ratio between epithelial basal cells and LCs were compared between ear and paw and showed no significance. Unpaired Student’s t-test (n=3 mice respectively). (e) *Hras^G12V/+^* mice have a higher epithelial basal cell density due to enhanced proliferation of epithelial stem cells *(Kl4-CreER; CX3CR1-GFP;Hras^G12V/+^)* and show consequent increase in immune cell densities in the basal layer of ear epidermal preps compared to Hras^+/+^ control mice *(Kl4-CreER; CX3CR1-GFP;Hras^+/+^;* representative images from 3 mice respectively). Epithelial nuclei are white (top panel). LCs are red and DETCs are green (bottom panel). Scale bar, 30 μm. (f-g) Cell number quantifications were compared between *Hras^+/+^* and Hras^*G12V/+*^ mice and found to be significant for all three cell types. Epithelial basal cell number (f) and LC and DETC number (g) in 0.25 mm^2^. * *p*<0.05 and ** *p*<0.01, unpaired Student’s t-test (n=3 mice respectively). (h) The ratios between epithelial basal cells and immune cells were compared between *Hras^+/+^* and *Hras^G12V/+^* mice separately and showed no significance when comparing the ratio of LCs, and appeared similar when comparing that of DETCs with slight statistical significance. * *p*<0.05, unpaired Student’s t-test (n=3 mice respectively). (i) *Cdkn1b;K14-rtTA* mice *(huLangerin-CreER; Rosa-stop-tdTomato; CX3CR1-GFP; K14-H2BCerulean; tetO-Cdkn1b; K14-rtTA)* have lower density due to blocked proliferation of epithelial cells and show corresponding lower density of immune cells in the basal layer of the epidermis of the ear compared to *Cdkn1b* control mice *(huLangerin-CreER; Rosa-stop-tdTomato; CX3CR1-GFP; Kl4-H2BCerulean; tetO-Cdkn1b)* at day 3 post induction with 1 mg/ml doxyclycine, 1 % sucrose in their drinking water (representative images from 3 mice respectively). Epithelial nuclei are white (top panel). LCs are red and DETCs are green (bottom panel). Scale bar, 30 μm. (j-k) Cell number quantifications were compared between *Cdkn1b* and *Cdkn1b;rtTA* mice and found to be significant for epithelial basal cells and LCs with a correlating trend for DETCs. Epithelial stem cell number (j) and LC and DETC number (k) in 0.25 mm^2^. * *p*<0.05 and ** *p*<0.01, unpaired Student’s t-test (n=3 mice respectively). (l). The ratios between epithelial basal cells and immune cells were compared between *Cdkn1b* and *Cdkn1b; K14-rtTA* mice separately and directly showed no significance for both LCs and DETCs. Unpaired Student’s t-test (n=3 mice respectively).

To test whether the ratio between epithelial basal cells and LCs and DETCs is regulated, we first genetically altered the density of LCs and DETCs. Specifically, we used an inducible diphtheria toxin model driven by the Langerin promoter to eliminate LCs *(Langerin-Diphtheria Toxin Receptor or Lang-DTR)* [44] and a null mouse model to eliminate DETCs *(TCRδ KO)* [45]. We found that the loss of either immune cell type did not affect the density of epithelial basal cells nor the epidermal thickness of both the ear and the paw epidermis. (**Supplementary Figure 1a-c,e, Supplementary Figure 2a-d**). Consistent with previous findings, we found that eliminating LCs had no effect on the density of DETCs in the ear, and vice versa [46–48] **(Supplementary Figure 1d, f)**; additionally, the remaining subtype maintained a largely constant ratio with epithelial basal cells **(Supplementary Figure 1g, h**). To probe this relationship in the context of depletion of both LCs and DETCs, we generated conditional mice in which both LCs and DETCs can be depleted simultaneously by inducible expression of diphtheria toxin *(huLangerin-CreER; TCR δ-CreER; Rosa-GFP-stop-DTA)* [35, 49, 50]. While LCs and DETCs together constitute a sizeable cellular fraction of the epidermis, their depletion did not significantly affect epithelial basal density or epidermal thickness **(Supplementary Figure 2e-g**). These data show that the densities of LCs and DETCs do not impact the density of epithelial basal cells or overall epidermal architecture.

To investigate the converse relationship of whether epithelial basal cell density regulates the density of LCs and DETCs, we altered the number of epithelial basal cells by either enhancing or blocking their proliferation. To increase epithelial basal cell density, we utilized a previously established genetic model with enhanced stem cell proliferation *(K14-CreER; Hras^G12V/+^)* [51, 52]. Remarkably, the density of both LCs and DETCs increased to match the higher epithelial basal cell number within six weeks after induction with tamoxifen, thereby maintaining similar epithelial-to-immune cell ratios observed during homeostasis in wild-type mice **(Figure 2e-h, Supplementary Figure 3a, Supplementary Figure 4a-j**).

In order to decrease epithelial basal density, we used mice with a tet-inducible *Cdkn1b* (or *p27)* overexpression allele in epithelial stem cells *(tetO-Cdkn1b; K14-rtTA)* [53, 54]. Doxycycline administration to the mice results in specific inhibition of basal stem cell proliferation and reduces their density in the epidermis within a day [38, 51, 55]. We observed an immediate decrease in both LC and DETC numbers leading to the maintenance of the ratios between epithelial basal cells and LCs or DETCs **(Figure 2i-l, Supplementary Figure 3b, c, Supplementary Figure 5a-d, Supplementary Figure 6a-h)**. Notably, we did notice a modest reduction in the number of all suprabasal cells at day 3 in the *Cdkn1b;rtTA* mice compared to controls **(Supplementary Figure 5e).** Overall, these data indicate a unidirectional relationship by which the number of epithelial cells dictates the number of LCs and DETCs, and where this regulation is rapid and robust.

Having demonstrated the positional stability and density regulation of LCs and DETCs, we next asked how their organization is orchestrated at the populational level. To this end, we used quantitative image analysis to examine the relative position of immune cells within the basal layer of the epidermis. We compared the experimental distributions of either LCs or DETCs to the artificially computer-generated random distributions of the same type of immune cell. In addition, we measured the distance to its nearest neighbors as well as the area occupied by each immune cell using Voronoi diagrams **(Figure 3a)**. Compared to the artificially generated random distributions, LCs maintained a larger average minimum distance from themselves, as did DETCs, which was further characterized by a tighter distribution around the mean (**Figure 3b**). This indicates that LCs and DETCs are each regularly and non-randomly distributed among themselves, consistent with observations for human LCs [56, 57].

**Figure 3.**
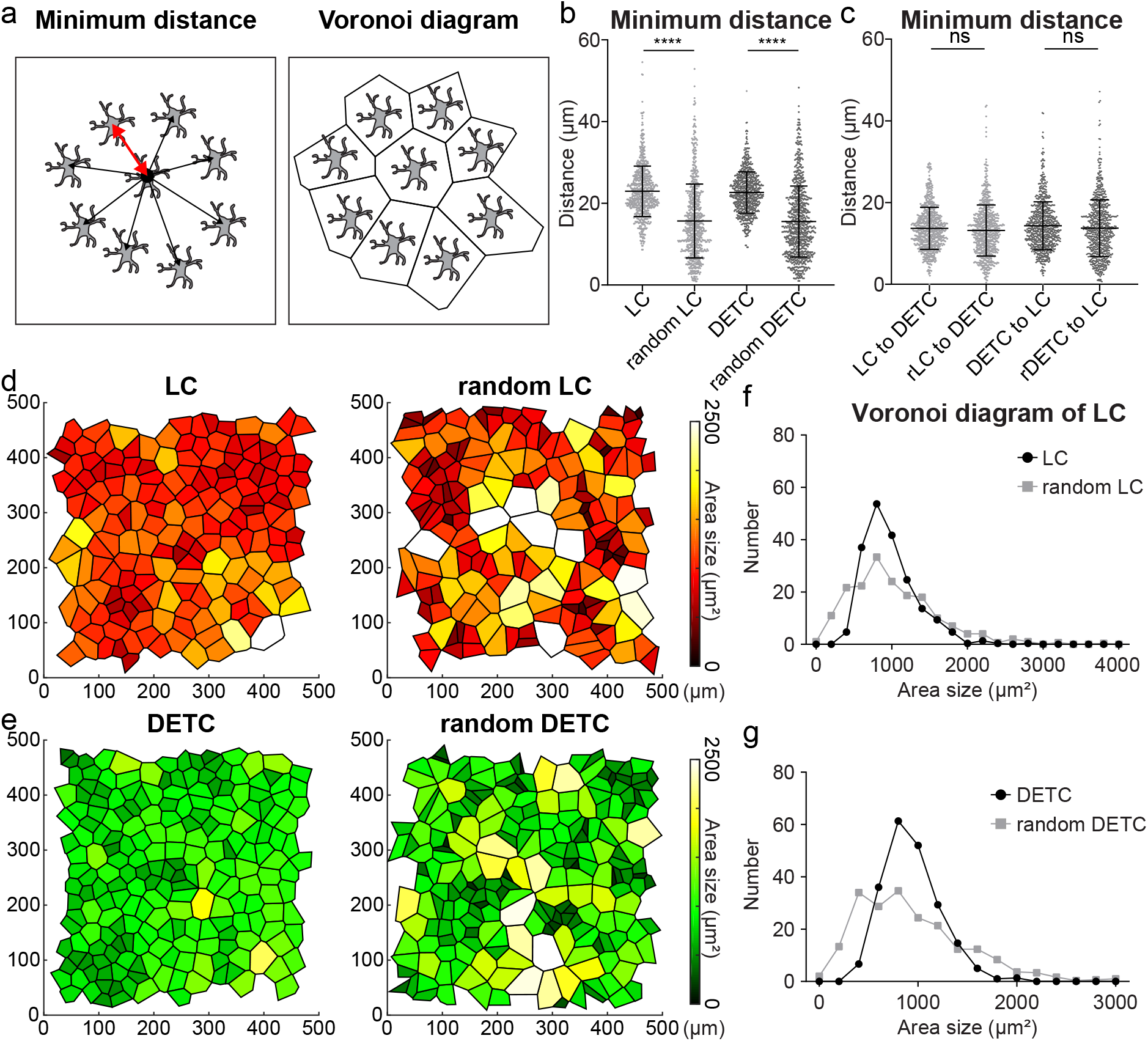
LCs and DETCs are organized in a tiling pattern. (a) Schematics of minimum distance and Voronoi diagram analyses used for quantification. (b) The minimum distance between the same type of immune cells (between LCs or between DETCs) was quantified as described in the schematic from experimental images and compared to random plot images, showing longer averages and significantly tighter distributions in both LC and DETC. **** *p*<0.0001, unpaired Student’s t-test (n=3 *huLangerin-CreER; Rosa-stop-tdTomato; CX3CR1-GFP; K14-H2BCerulean* mice respectively). (c) The minimum distance between different types of immune cells (between LCs and DETCs) was quantified from experimental images and compared to random plot images and showed no significant difference. Unpaired Student’s t-test (rLC, random LC; rDETC, random DETC; n=3 mice respectively). (d-e) Voronoi diagrams were generated from the experimental images (left) and the random plots (right) to represent the area occupied by each immune cell. The experimental images show a more regular distribution than the random plots. LC and random LC (d). DETC and random DETC (e). Area size 0.25 mm^2^. Polygons are color coded by size as indicated in the legend (representative images from 3 mice). (f, g) Quantification of area size from the Voronoi diagram between the experimental images (black) and random plots (gray). The experimental images show tighter distributions than the random plots. LC and random LC (f). DETC and random DETC (g) (n=3 mice respectively).

To determine whether the distribution of LCs is dependent on the distribution of DETCs and vice versa, we compared actual imaging coordinates of one immune population to actual or artificially generated random distributions of the other. Through this analysis, we found that LCs and DETCs do not maintain a specific distance from each other **(Figure 3c**). This corroborates the above findings that the depletion of LCs or DETCs does not impact the density or distribution of the remaining population **(Supplementary Figure 1a, Supplementary Figure 7a-d**).

To further understand the global distribution of immune cells, we measured the area that each LC and DETC occupies using a Voronoi diagram. We found that the Voronoi diagrams of both LCs and DETCs revealed a more uniform and equal partitioning than those created using a set of artificially generated randomly positioned cells, reinforcing the regularity of these immune cell patterns (**Figure 3d-g**). Together, these data indicate that LCs and DETCs are not randomly distributed in the epidermis and that the tiling patterns of LCs and DETCs are independently regulated.

Our data show that LCs and DETCs have a relatively stable position and are regularly distributed during homeostasis of the regenerative epidermis. Based on these findings, we questioned how the loss of either LCs or DETCs would affect their respective patterns in the days following this loss. Given that newly generated cells would arrive only weeks later [33, 44, 58–61], we hypothesized three scenarios: 1) the void left upon cell loss would persist because surviving LCs and DETCs maintain their initial positions, 2) the void would be filled by remaining LCs or DETCs to reestablish a pattern, or 3) the void would be filled by remaining LCs or DETCs without reestablishing a pattern. In order to explore these possibilities, we used two complementary approaches to eliminate LCs or DETCs and examined the response of surviving neighboring immune cells.

First, we used our previously established single cell laser ablation to precisely eliminate a small population of LCs [38, 62] and tracked the surviving epidermal neighboring LCs over the next few days. Strikingly, we found that LCs moved into the ablated region and established a regular pattern within the previously ablated regions **(Figure 4a, b**). Moreover, using the same approach we found that DETCs behaved similarly (**Supplementary Figure 8a, b)**.

**Figure 4.**
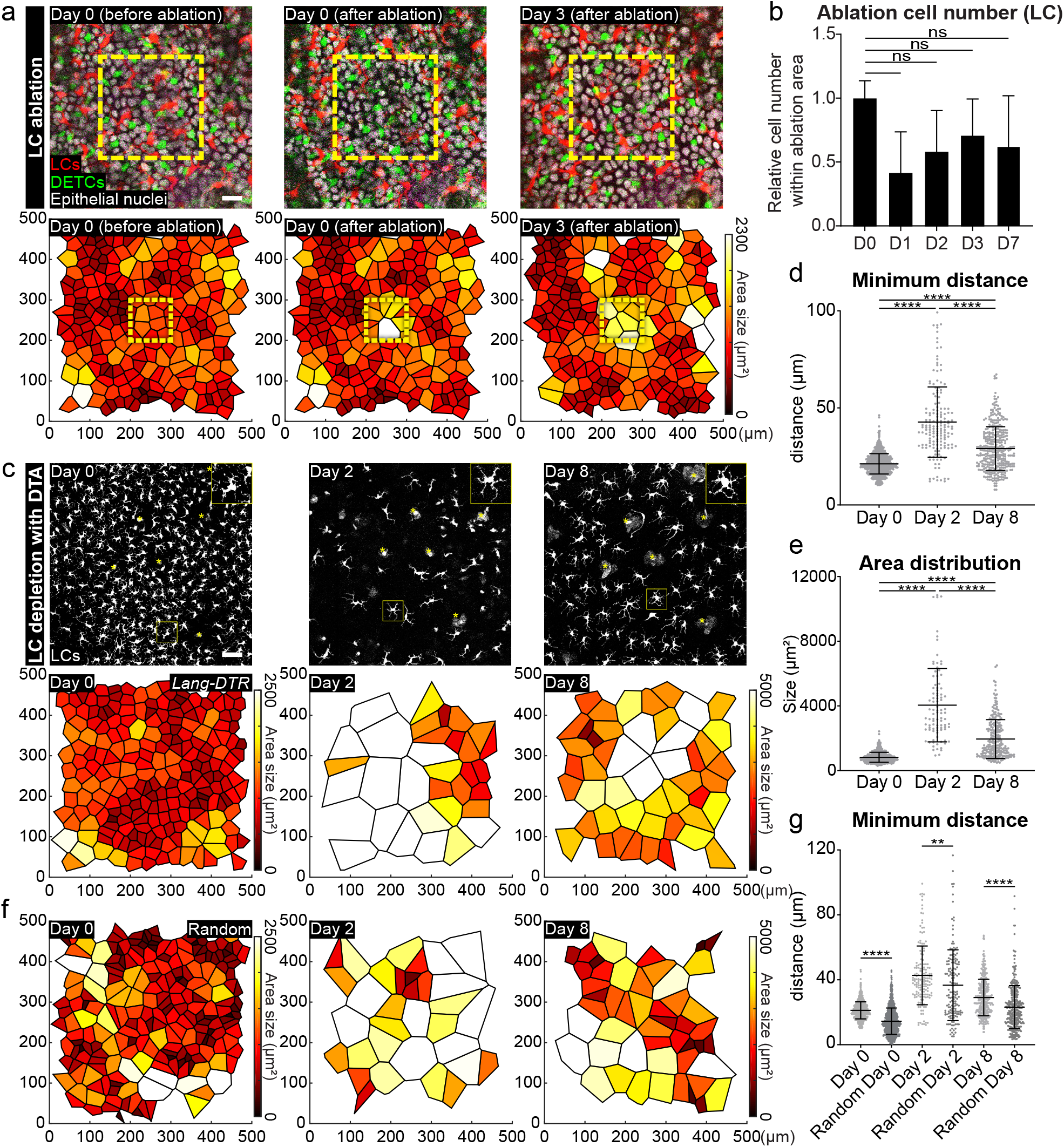
Surviving LCs re-establish a regular pattern after neighboring LC loss. (a) Local laser ablation of LCs. LCs within the yellow box (100 μm X 100 μm) are ablated by multiphoton laser (810 nm) and the same region is revisited 3 days after the ablation. The upper panel shows actual images from a mouse from day 0 before ablation to day 3 post ablation. Scale bar, 10μm. The lower panel displays the Voronoi diagram for LCs generated from the images at each timepoint and encompasses a larger area around the ablation site. Neighboring LCs migrate into the ablated region and re-establish a regular pattern. Area size 0.25 mm^2^ (representative images from 4 mice). (b) The LC pattern within the ablated region was quantified from day 1 to day 7 and compared to the initial number at day 0. Unpaired Student’s *t*-test (n=4 mice respectively). (c) Global depletion of LCs. One dose of diphtheria toxin (2 ng/g of body weight) was injected intraperitoneally (i.p.) into *Lang-EGFP; Lang-DTR (Lang-DTR)* mice for the acute depletion of LCs. The same region was revisited at day 2 post injection when 82 % of LCs have died and shows uneven pattern similar to random. The regular pattern is recovered at day 8 with fewer cells (n=3 mice respectively). The upper panel shows LCs in the basal layer at each imaged timepoint. Scale bar, 50μm. Yellow asterisk denotes hair follicles. The lower panel is the Voronoi diagram for LCs generated from the images at each timepoint. (d) The minimum distance between LCs at each timepoint (day 0, 2 and 8) were quantified. The minimum distance is increased with broader distribution right after the depletion (day 2) but recovers at day 8. **** *p*<0.0001, unpaired Student’s *t*-test (n=3 mice respectively). (e) Quantification of area size from the Voronoi diagram from LCs at each timepoint. Similar to the minimum distance, the area distribution is also recovers at day 8. **** *p*<0.0001, unpaired Student’s *t*-test (n=3 mice respectively). For Voronoi diagrams, polygons are color coded by size (as indicated in the legend). (f) Voronoi diagrams generated from equal numbers of artificially generated random LC positions. The patterns show greater variability in polygon size compared to experimental images at both day 0 (before the depletion) and day 8 (after the recovery) (representative images from 3 random plots). (g) The same minimum distance plots from (d) were compared to equal numbers of artificially generated random positions. The experimental groups show longer minimum distances with tighter distributions than random plots at all time points. Right after depletion (day 2) shows a smaller significance. \*\**p*<0.01 and **** *p*<0.0001, unpaired Student’s *t*-test (n=3 mice respectively).

To complement these findings, we also assessed how the loss of a much larger number of cells affects immune cell distribution by treating *Lang-DTR* mice with an acute dose of diphtheria toxin. Tracking of the same epidermal areas over time revealed gaps in the immune cell pattern at day 2 after treatment, and LCs did not recover in number by day 8 **(Figure 4c top panel, Supplementary Figure 9a)**[55]. Importantly, the surviving LCs in the diptheria toxin-treated *Lang-DTR* mice migrated and reestablished a non-random pattern in the skin basal layer **(Figure 4c bottom panel, d, e)**. We also compared the minimum distance of LCs at each of the timepoints to those of random distributions and found that at day 8, the LCs had much less variation in their minimum distances to each other compared to artificially generated random distributions than at day 2 suggesting that they were actively recovering their pattern **(Figure 4f, g, Supplementary Figure 9b)**. Taken together, these results indicate that the loss of LCs and DETCs during homeostasis triggers a response from surviving, neighboring immune cells to move into voided space and re-establish a homeostatic tiling pattern in the epidermis of adult mice.

To define mechanisms underlying how LCs and DETCs actively maintain their tiling patterns and minimum distance, we sought to target a regulator of dendrite development, the actin remodeling molecule Rac1, a member of the Rho GTPase family [63]. In neuronal self-avoidance, dendrites arising from the same neuron do not overlap [64, 65]. This striking parallel with LCs led us to hypothesize that dendrites may play a role in establishing their tiling pattern. Conditional loss of function of Rac1 in LCs *(Rac1^fl/fl^* [66]; *huLangerin-CreER* [35] treated with tamoxifen, referred to as LC^Rac1KO^) resulted in a lack of LC dendritic radial organization and fewer dendritic branches, consistent with previous studies in neurons and dendritic cells [63, 67, 68], as well as reduced LC density which may be due to an initial increase in migration out of the epidermis [69] (**Figure 5a; Supplementary Figure 9a, Supplementary Figure 10a-d**).

**Figure 5.**
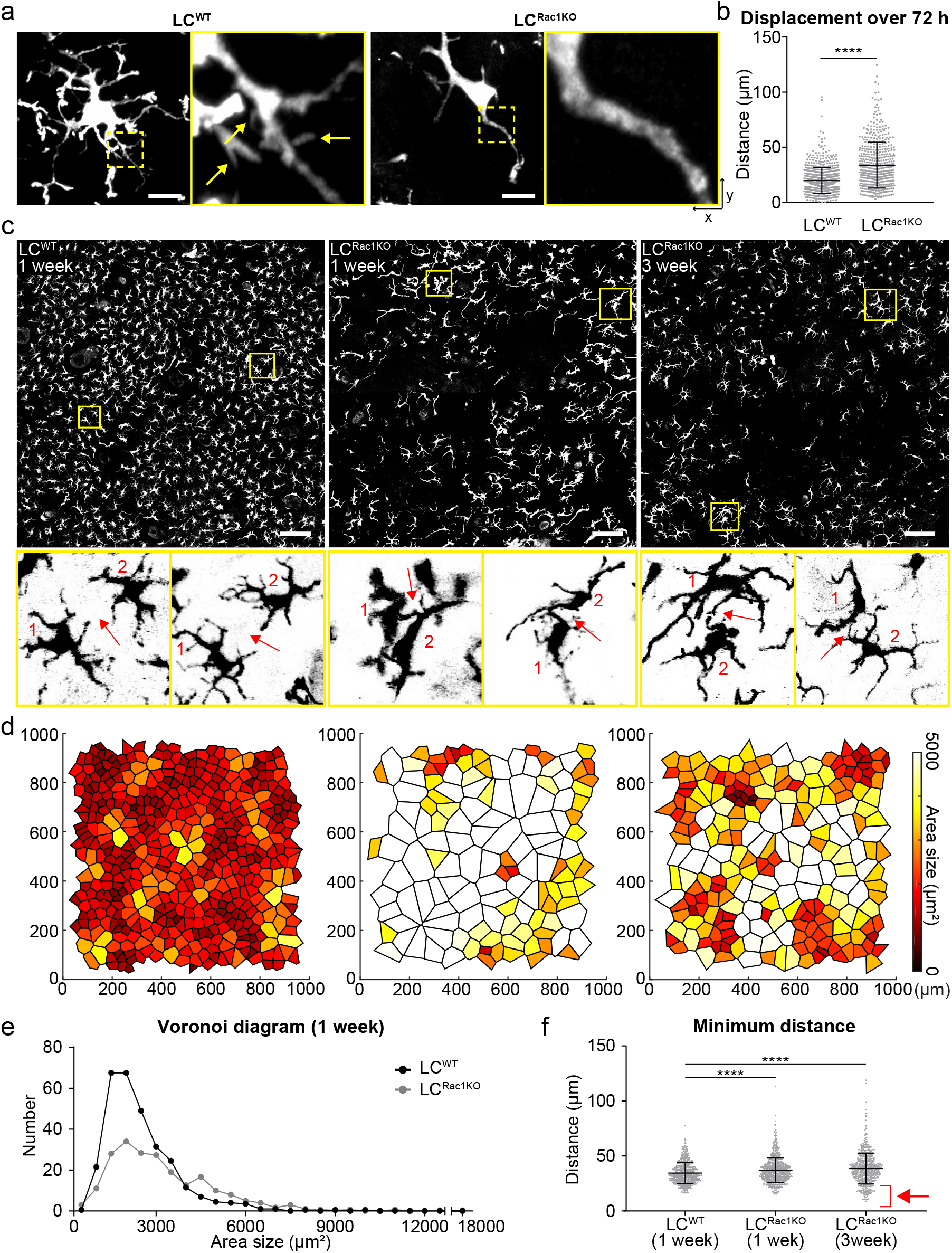
Rac1 deficiency in LCs causes altered dendritic morphology, higher mobility, and loss of patterning. (a) Rac1 is eliminated specifically in LCs by treating *huLangerin-CreER; Rosa-stop-tdTomato; Rac1^fl/fl^* mice with 2 mgs of tamoxifen on 5 consecutive days. Rac1 KO LCs (LC^Rac1KO^) show lack of dendritic radial morphology and fewer dendrites when compared to the control LCs (LC^WT^) by Airyscan (n=3 mice respectively). Scale bar, 10μm. (b) The displacement of LC^Rac1KO^ was found to be larger than that of LC^WT^ over 72h in revisit analysis showing that the disrupted pattern is not due to the inability of LC^Rac1KO^ to migrate,. Area quantified 0.56 mm^2^. **** *p*<0.0001, unpaired Student’s *t*-test (n=3 mice respectively). (c) There is a disruption of the regular pattern in LC^Rac1KO^ compared to LC^WT^ 1 week after tamoxifen injection (middle panel) which persists at 3 weeks after tamoxifen injection (right panel; n=2 control mice, n=3 Rac1 KO mice). The top panel shows LCs in the basal layer (scale bar, 100 μm) and the bottom panel shows insets highlighting the distance between neighboring LCs (red arrows). Despite the decrease in density, LC^Rac1KO^ are in much closer contact than LC^WT^. (d) The Voronoi diagram for LCs generated from the images in the top panel of (c). (e) Quantification of area size from Voronoi diagrams of LC^WT^ and LC^Rac1KO^ 1 week after tamoxifen injection. LC^Rac1KO^ show larger partition areas than LC^WT^ (n=3 mice respectively). (f) Quantification of minimum distance comparing LC^Rac1KO^ at week 1 and week 3 to LC^WT^ shows that LCRac1^KO^ lose the ability to maintain a consistent minimum distance with their neighbors (red arrow). **** *p*<0.0001, unpaired Student’s *t*-test (n=3 mice respectively). For Voronoi diagrams, polygons are color coded by size (as indicated in the legend).

To investigate how Rac deletion affects positional stability, we performed timelapse imaging over a few hours and measured the relative LC^Rac1KO^ displacement over 72 hours [55, 69]. Intriguingly, while LC^Rac1KO^ had similar slow migratory speeds as LC^WT^ (4 h; **Supplementary Movie 3**), they showed greater displacement when compared to WT or genetic models with comparable reduced LC density (72 h; **Figure 5b, Supplementary Figure 9c)**. These results indicate that a conditional loss of Rac1 function in LCs does not compromise their migration but disrupts their apparently stable positioning in the epidermis. These data suggest that Rac1, and potentially dendrites, are important to constrain the movement of LCs in the epidermis.

In some ways, these cellular phenotypes resemble an LC activation state in terms of their changes in morphology and decrease in density [31, 70, 71]. However, LC^Rac1KO^ lacked key molecular markers of activation, such as the upregulation of MHCII or CD86 when compared to LC^WT^ or physiologically activated LCs from lymph nodes **(Supplementary Figure 11a-e)**. In addition, LCs maintained expression of the adherent junction component E-cadherin **(Supplementary Figure 11f-h)**, which is downregulated during LC activation [72, 73]. These data suggest that LC^Rac1KO^ in the epidermis are not activated and maintain cell-cell contact with epithelial cells.

To assess LC^Rac1KO^ immune cell patterning, we revisited the same region over time. Strikingly, LC^Rac1KO^ lost their tiling pattern by week 1 post induction and this irregular distribution persisted for at least 3 weeks (**Figure 5c-e**). For instance, LC^Rac1KO^ cells at 3 weeks have a wider range of minimum distances and come into closer contact than LC^WT^ cells, consistent with a loss of self-avoidance **(Figure 5c bottom panel, f)**. The altered minimum distance and increased displacement of LC^Rac1KO^ does not appear to be the result of lower LC numbers as other models with comparable drops in LC density such as the *Cdkn1b;rtTA* and *Lang-DTR* models still maintained relatively stable positions and non-random patterns (**Supplementary Figure 9b, c**). Collectively, these data suggest that Rac1 ablation in LCs results in dendrite defects, an inability to remain spatially stable, and failure to maintain their non-random tiling pattern.

This study set out to investigate population dynamics between immune cells and epithelial cells in the context of adult skin homeostasis. We discovered previously unknown principles and behaviors that maintain immune cell density and spatial coverage during homeostasis. Specifically, immune cells in the epidermis maintain a relatively stable position in the face of perpetual changes in their epithelial neighborhood. At the same time, immune cells exhibit dynamic behaviors, continuously exploring epithelial cell junctional spaces through their rapid dendritic movements. We identified two distinct levels of control of the immune cell population. On the one hand, we found that immune cell populations modulate their densities based on the density of their epithelial cell neighbors, so that a constant epithelial:immune cell ratio is maintained across the basal layer of the epidermis. On the other hand, we uncovered that both LCs and DETCs are organized in a tiling pattern that is actively maintained regardless of their density in the epidermis, but which relies on the positional stability associated with Rac1 **(Figure 6)**. Taken together, our results identify novel relationship dynamics between immune cells and epidermal cells, and provide insight into how immune cells organize and adapt to local changes during adult homeostasis. We propose that dynamic regulation within and between the diverse cell types of the skin maintain global organ homeostasis and enable adaptations to local changes in the environment during adult homeostasis.

**Figure 6.**
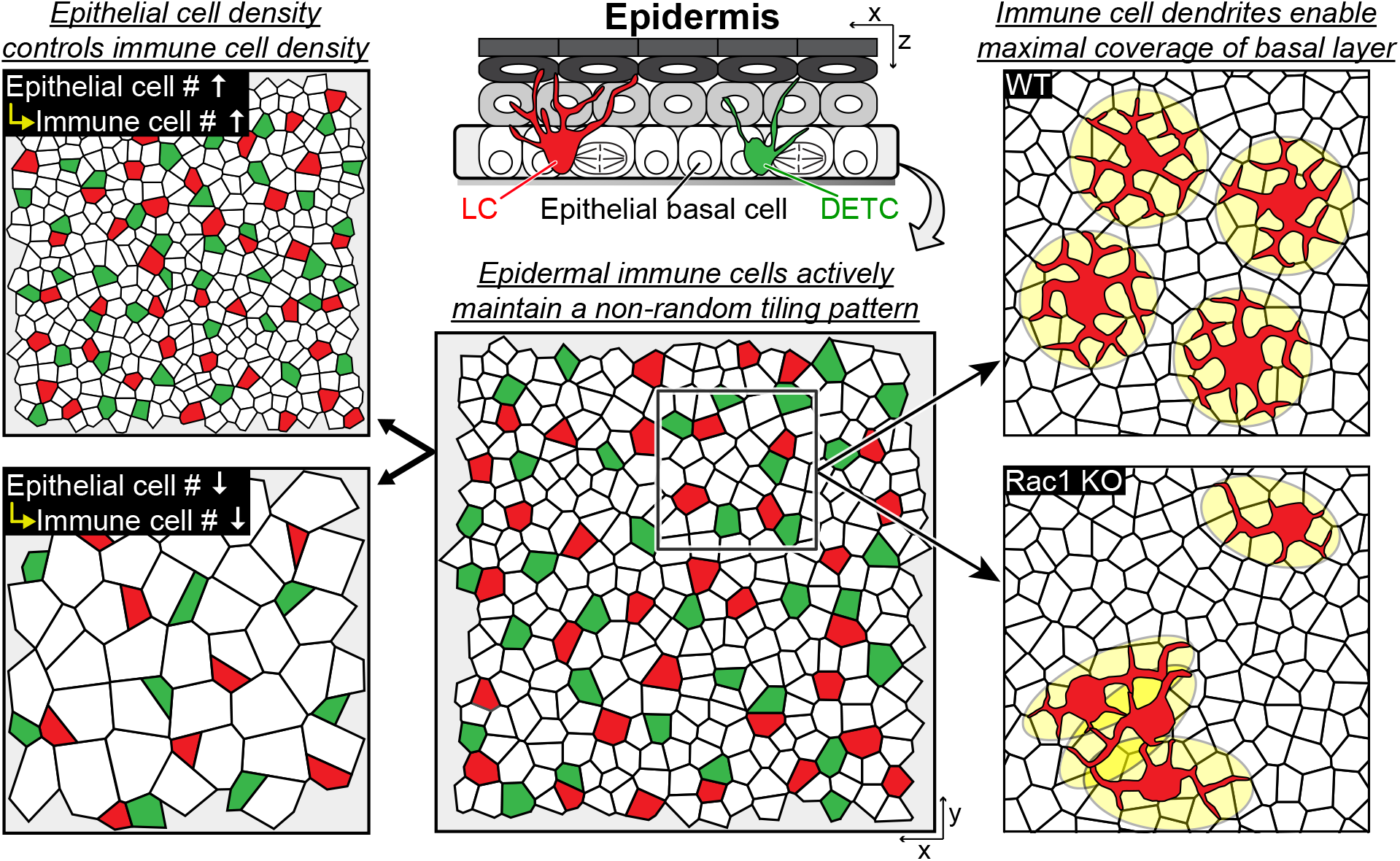
Skin-resident immune cells actively coordinate their distribution with epithelial basal cells. During homeostasis, LCs and DETCs, Langerhans cells (LCs) and dendritic epidermal T cells (DETCs), actively maintain a non-random distribution. Although these immune cells continuously move within epithelial basal cell neighbors, they sustain their regular tiling pattern. Moreover, epithelial stem cells determine the density of LCs and DETCs in the skin epidermis. However, LCs and DETCs do not impact the density of epithelial stem cells nor the architecture of the epidermis signifying a unidirectional regulation of density. Finally, immune cell dendrites enable maximal coverage of the basal layer and do not come into contact with neighboring immune cells in homeostasis and their dendritic behaviors, mediated by Rac1, coordinate their patterned organization.

## Methods

### Mice and experimental conditions

*huLangerin-CreER* [35], *Rosa-stop-tdTomato* [36], *CX3CR1-GFP* [37], *K14-CreER* [74], *K14-rtTA* [54], *tetO-Cdkn1b* [53], *Lang-DTR* [44], *Rac1^fl^* [66], *Rosa26 LSL H2B mCherry* [75], *TCRδ-CreER [49]* and *Rosa-GFP-stop-DTA* [50] mice were obtained from Jackson Laboratories. *K14-H2BCerulean* mice were generated by the Yale Transgenic Facility. *Lang-EGFP* [44] mice were obtained from N. Anandasabapathy (Brigham & Women’s Hospital), *Hras^G12V^* [76] mice were obtained from Slobodan Beronja (Fred Hutch) and *TCRδ KO* [45] mice were obtained from Akiko Iwasaki (Yale University. To visualize LCs, DETCs and epithelial cells at the same time, *huLangerin-CreER; Rosa-stop-tdTomato; CX3CR1-GFP; K14-H2BCerulean* mice were generated and Cre expression was induced with a single intraperitoneal injection of tamoxifen (2 mg in corn oil). To block the proliferation of epithelial cells, the *huLangerin-CreER; Rosa-stop-tdTomato; CX3CR1-GFP; K14-H2BCerulean* mice were mated with *K14-rtTA* and *tetO-Cdkn1b (huLangerin-CreER; Rosa-stop-tdTomato; CX3CR1-GFP; K14-H2BCerulean; tetO-Cdkn1b; K14-rtTA)* and these mice were given a single intraperitoneal injection of tamoxifen (2 mg in corn oil) as well as doxycycline (1 mg/ml) in potable water with 1% sucrose [55]. Doxycycline treatment was sustained until imaging was performed. Siblings without the *K14-rtTA* allele were used as controls *(huLangerin-CreER; Rosa-stop-tdTomato; CX3CR1-GFP; K14-H2BCerulean; tetO-Cdkn1b).* To enhance the proliferation of epithelial cells *K14-CreER;CX3CR1-GFP* or *K14-CreER; Rosa26 LSL H2B mCherry* mice were mated with *Hras^G12V^* mice *(K14-CreER; CX3CR1-GFP; Hras^G12V/+^* or *K14-CreER; Rosa26 LSL H2B mCherry; Hras^G12V/+^* respectively) and Cre expression was induced with a single intraperitoneal injection of tamoxifen (2mgs in corn oil) [51]. Wild type were siblings used as a control *(K14-CreER; CX3CR1-GFP; Hras^+/+^; K14-CreER; Rosa26LSL H2B mCherry;Hras^+/+^*). To visualize and deplete LCs, *Lang-EGFP* mice were mated with *Lang-DTR (Langerin-Diphtheria Toxin Receptor)* mice in order to boost the weak EGFP signal of the *Lang-DTR* mice *(Lang-EGFP; Lang-DTR).* These mice were given either a single intraperitoneal injection of diphtheria toxin (2 ng/g of body weight in PBS) at postnatal day 21 for the partial depletion of LCs or 1μg/body weight for full depletion depending on the experimental conditions [44, 77]. To knock-out Rac1 in the LCs, *huLangerin-CreER; Rosa-stop-tdTomato; CX3CR1-GFP; K14-H2BCerulean* mice were mated with Rac1^fl/fl^ mice *(huLangerin-CreER; Rosa-stop-tdTomato; CX3CR1-GFP; K14-H2BCerulean; Rac1^fl/fl^)* and these mice were given five intraperitoneal injections of tamoxifen (2 mg in corn oil per day for 5 days). Wild type siblings used as controls *(huLangerin-CreER; Rosa-stop-tdTomato; CX3CR1-GFP; K14-H2BCerulean; Rac1^+/+^).* To deplete LCs and DETCs simultaneously, we crossed *Rosa-GFP-stop-DTA* to *huLangerin-CreER; TCRδ-CreER* mice and gave them 2 intraperitoneal injections of tamoxifen (2mgs in corn oil for 2 consecutive days). Wild type siblings used as controls *(Rosa-GFP-stop-DTA).* Mice from experimental and control groups were randomly selected of either gender for live imaging experiments. No blinding was done. All procedures involving animal subjects were performed under the approval of the Institutional Animal Care and Use Committee (IACUC) of the Yale School of Medicine.

### *In vivo* imaging

Mice were anesthetized using an isofluorane chamber and anesthesia was maintained throughout the course of the experiment with vaporized isofluorane delivered by a nose cone as previously described [40]. Image stacks were acquired with a LaVision TriM Scope II (LaVision Biotec, Germany) laser scanning microscope equipped with a tunable Two-photon Vision II Ti:Sapphire (Coherent, USA) Ti:Sapphire laser and tunable Two-photon Chameleon Discovery Ti:Sapphire larer (Coherent,USA). To acquire serial optical sections, a laser beam (940 nm for *Lang-EGFP*, *Lang-DTR*; 965 nm for *huLangerin-CreER; Rosa-stop-tdTomato; CX3CR1-GFP; K14-H2BCerulean* mice; 940 nm, 1040nm and 1120nm for wholemount staining) was focused through a 20X or 40X water immersion lens (N.A. 1.0 and 1.1 respectively; Zeiss, USA) and scanned with a field of view of 0.5 × 0.5 mm^2^ or 0.25 × 0.25 mm^2^ respectively at 600 Hz or through a 25X water immersion lens (N.A. 1.0; Nikon, Japan) and scanned with a field of view of 0.44 × 0.44 mm^2^ at 600 Hz. Z-stacks were acquired in 1-3 μm steps to image a total depth of 200 μm of tissue. To visualize large areas, 3-36 tiles of optical fields were imaged using a motorized stage to automatically acquire sequential fields of view as previously described [40]. Visualization of collagen was achieved via second harmonic signal (SHG) using blue channel at 940 nm imaging wavelength. For time-lapse imaging, serial optical sections were obtained in a range of 1 to 12 minute intervals depending on the experimental setup. The duration of time-lapse imaging ranged from 1-4 hours. Frequent daily imaging can cause a drop in cell density in epithelial basal cells, LCs and DETCs, and experiments were either normalized to the control or repeated with single timepoints to confirm findings when necessary.

### Laser Ablations

Single cells were ablated with a tunable Two-photon Vision II Ti:Sapphire (Coherent, USA) Ti:Sapphire laser (810nm wavelength, with 15-20 % power, PMTs at 100 %) for 3 seconds using a “live” scan focused through a 63X water immersion lense (N.A. 1.5, Zeiss USA) at 800Hz and within a 1 um^2^ size window in the center of the cell body.

### Image Analysis

Raw image stacks were imported into Fiji (NIH, USA) or Imaris software (Bitplane/Oxford Instruments) for further analysis. Imaris software was used to track cells and obtain xyz coordinates from individual tracked cells over time. All cell tracks were individually examined and only cells that could be tracked for the full duration of a movie or of revisit images were included in the final analysis. Prism software (Graphpad) was used to graph the data. The tiled images were stitched by a grid/collection stitching plugin in Fiji. Migrating cell tracking for rose plot analysis and 3D reconstitution were performed in Imaris software. Voronois, displacement analysis and minimum distances were analyzed using Matlab (Mathworks, USA).

### Whole mount staining

Ear tissues were processed for whole mount staining. Briefly, ears were incubated epidermis side up in 5 mg/ml dispase II solution (Sigma, 4942078001) at 37 °C for 15 minutes and epidermis was removed from the dermis. The epidermis was fixed in 4 % paraformaldehyde in PBS for 10 minutes at room temperature, washed in PBS, and then blocked with 0.2 % Triton, 5 % NDS, 1 % BSA in PBS. The samples were then incubated in primary antibodies for 24 hours and secondary antibodies for ~1 hour at room temperature. Primary antibodies used were as follows: purified Armenian Hamster anti-mouse T cell receptor γ/δ (BioLegend, 118101), purified rat anti-mouse MHC class II or MHC class II Alexa 488 (BioLegend, 107602,107616), purified rat anti-mouse CD45 (Biolegend, 103102), purified mouse anti mouse/human CD207 (Biolegend, 144202), rabbit anti-mouse/human Ki67 (abcam, ab15580), rabbit antimouse Phospho-PI3 Kinase p85 (Tyr458)/p55 (Tyr199) (Cell Signaling, 4228S), and Cleaved-Caspase 3 (Asp175) (5A1E) Rabbit mAb (Cell Signaling, 9664). Secondary antibody used were as follows: Goat anti-hamster Alexa Fluor 488, Goat anti-hamster Alexa Fluor 568, Goat anti-rat Alexa Fluor 488, Goat anti-rat Alexa 568, Goat-anti rabbit Alexa Fluor 568 and Goat-anti rabbit Alexa 633 (ThermoFisher). Tissue was then incubated with Hoechst 33342 (H3570, Becton Dickinson) for 15 minutes, washed with PBS and mounted on a slide or directly mounted on a slide with DAPI (Vector; H1200) and imaged on a LaVision TriM Scope II as described in *in vivo* imaging. For Airyscan imaging, ear tissue was dissected and fixed in 4 % paraformaldehyde in PBS for 1 hour at 4 °C. Fixed tissue was washed in PBS for one hour at room temperature on a rocking platform, then mounted on a slide with Vectashield Anti-fade mounting medium (Vector Laboratories) with a #1.5 coverslip. Airyscan imaging was performed on a Zeiss LSM880 with 488 nm and 568 nm laser lines and rendering was done within the Zeiss acquisition software at default settings.

### Flow Cytometry

*huLangerin-CreER; Rac1^+/+^* (n=4) and *huLangerin-CreER; Rac^fl/fl^* (n=5) mice were euthanized for FACS two days after receiving five intraperitoneal injections of tamoxifen (2 mg in corn oil per day for 5 days) to reflect conditions utilized in imaging studies. *Cdkn1b* (n=3) and *Cdkn1b;rtTA* (n=3) mice were euthanized for FACS 3 days after the induction with doxycycline (1 mg/ml) in potable water with 1% sucrose. Epidermal and lymph node single cell suspensions were prepared for flow cytometry with a protocol adapted from previously described [71]. Briefly, single-cell suspensions of epidermal cells were obtained from ear skin and incubated for 1 hour at 37 °C in 0.3 % trypsin (Sigma-Aldrich) in 150 mM NaCl, 0.5 mM KCl and 0.5 mM glucose. The epidermis was physically separated from the dermis, minced, and the resulting cells were crushed and filtered through a 70 μm filter. Lymph nodes were incubated in 400 U/ml collagenase D (Roche Applied Science) for 30 min before filtration through a 70 μm filter. All samples were pretreated with rat serum (Sigma-Aldrich) and incubated with anti-mouse/human CD207 PE (Biolegend, 144204) and either anti-mouse MHCII APC (Biolegend, 107614), anti-mouse CD86 APC (Biolegend, 105012), or anti-mouse CD324 biotin (E-cadherin) (Thermofisher, 13-3249-82) at 4 °C. Biotin samples were incubated with secondary antibody Streptavidin-APC (Thermofisher, SA1005) for 20 minutes. Samples were run on a Becton Dickinson LSRII outfitted with Diva software and the data was analyzed using Flowjo v10.6.2.

### Statistics and reproducibility

Data are expressed either as absolute numbers or percentages ± S.D. An unpaired Student’s *t*-test was used to analyze data sets with two groups and * *p*<0.05, ** *p*<0.01, *** *p*<0.001 and **** *p*<0.0001 indicate a significant difference. Statistical calculations were performed using the Prism software package (GraphPad, USA). No statistical method was used to pre-determine sample size. Panels showing representative images are representative of at least two independent experiments and up to five, as indicated in the figure legends.

### Code availability

X-y Positions of immune cells were identified using Fiji. The minimum distance and displacement analyses were performed using the Matlab function “squareform”. The Voronoi tessellation used to determine nuclei neighbor relationships was performed using Matlab function “voronoi”. To make an artificially generated random pattern, random x-y positions were generated by Matlab function “randi”.

### Data availability

All data that support the conclusions are available from the authors on request.

## Acknowledgements

We thank A. Anderson for critical feedback on the manuscript. We thank N. Anandasabapathy for the *Lang-EGFP* mice, S. Beronja for the *Hras^G12V^* mice, and Akiko Iwasaki for the *TCRδ KO* mice. The work is supported by the HHMI Scholar award, NIH, grants no. 1R01AR072668-01. S.P was supported by The New York Stem Cell Foundation (NYSCF-D-F58). Y.B. is supported by the Centre National de la recherche Scientifique (CNRS), The Institut Curie, and the Institut National de la santé et de la recherche médicale (INSERM). E.M. was supported in part by the National Institute of Health (T32 GM007499). D.M. was supported by The National Insitute of Health (T32-GM007223-44).

## Author Contributions

S.P., C.M. and V.G designed experiments and analyzed data. S.P. performed 2-photon imaging, laser ablations, Matlab and IMARIS analysis, mouse genetics, and toxin injections. C.M. performed 2-photon imaging, whole mount staining, FACS preparatory work and analysis and mouse genetics. D.G. assisted in 2-photon imaging, Matlab and IMARIS analysis, and experimental discussions throughout this project. E.L. assisted with whole mounts and mouse genetics, Y.B. assisted with immune cell modeling and critical feedback of the manuscript. J.B. assisted with mouse genetics. C.P. assisted with the Hras model development. E.M. assisted with laser ablations and Matlab analysis. J.M. assisted with analysis and critical feedback on the manuscript. D.M. assisted with airy scan microscopy and critical feedback on the manuscript. A.S. assisted with quantifications of epithelial density and cell-cell contact. K.C. assisted with critical feedback on the manuscript.

## Competing Financial Interests

The authors declare no competing financial interests.

## Supplementary Figure Legends

**Supplementary Figure 1.**
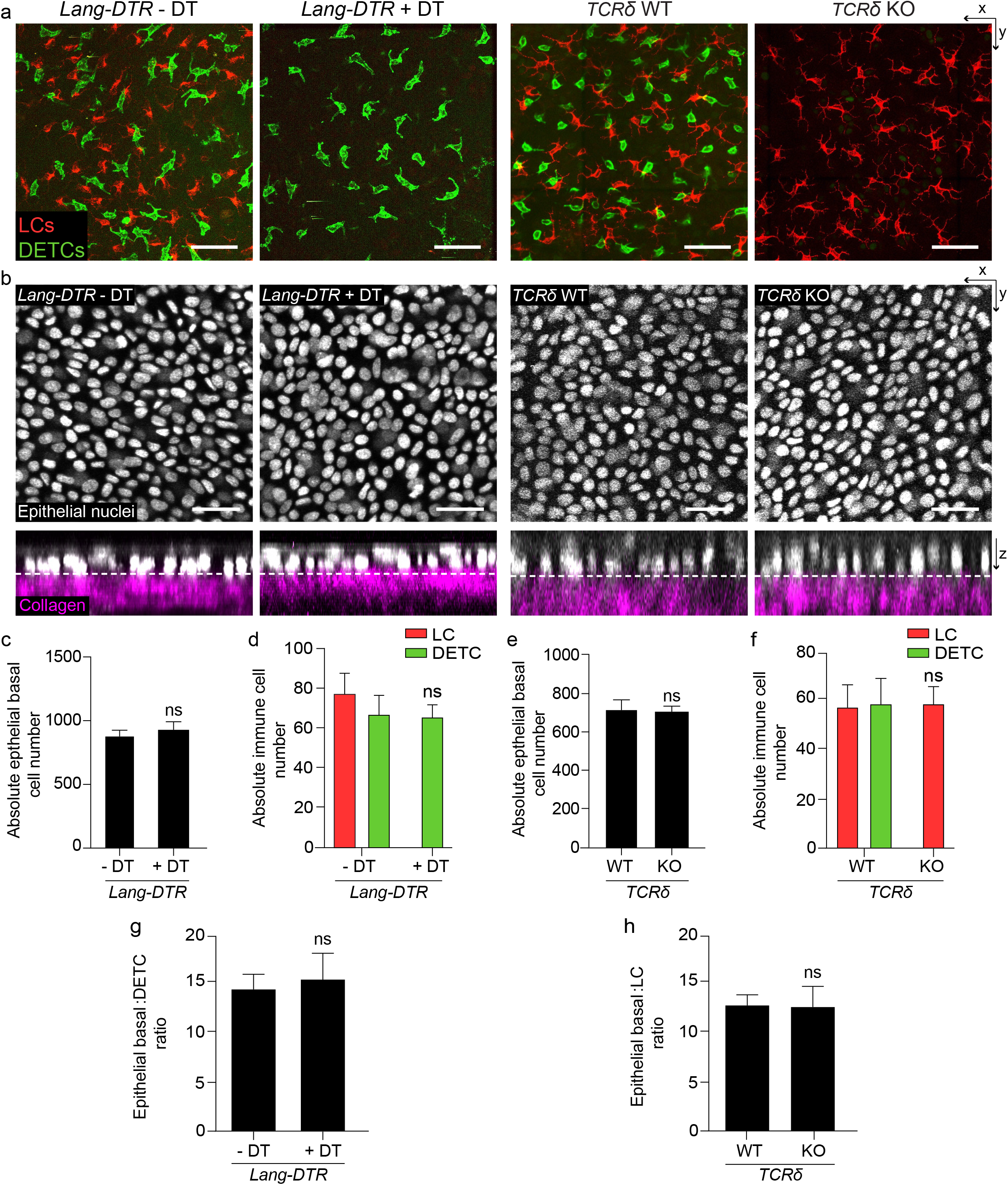
The depletion of LCs or DETCs does not impact epithelial basal density or epidermal architecture of the ear. (a) X-y view of the basal layer of the epidermis with LCs in red (anti-MHCII) and DETCs in green (anti-TCRγδ) comparing *Lang-DTR* control mice without the addition of diphtheria toxin (or - DT) to *Lang-DTR* mice given one dose of 1μg/body weight diphtheria toxin for the acute depletion of LCs 5 days before harvest (+ DT) (left panels) or comparing *TCδ WT* to *TCRδ KO* mice (right panels) (representative images from 3 mice each). The images show that the depletion of one immune population does not affect the density of the other. Scale bar, 50 μm. (b) X-y view of the basal layer and x-z view of the epidermis with epithelial nuclei in white and collagen in magenta, comparing *Lang-DTR* mice either - DT or + DT (left panels) or comparing *TCRδ WT* to *TCRδ KO* mice (right panels) (representative images from 3 mice each). The images show that the depletion of LCs and DETCs does not impact the density or epidermal architecture of epithelial stem cells. Scale bar, 30 μm. (c,e) Quantification of epithelial basal cell number (n=3 mice respectively) comparing *Lang-DTR* mice either - DT or + DT mice (c) and TCRδ WT to *TCRδ KO* mice (e). Unpaired Student’s *t*-test. (d,f) Quantification of LC and DETC cell numbers (n=3 mice respectively) comparing *Lang-DTR* mice either - DT or + DT, (d) and *TCRδ WT* to *TCRδ KO* mice (f). Unpaired Student’s *t*-test. (g,h) Ratio between epithelial basal and DETCs in *Lang-DTR*+ DT (g) and LCs in *TCRδ KO* (h) compared to their respective control mice (n=3 mice respectively). Area quantified for cell number 0.0625 mm^2^. Unpaired Student’s *t*-test.

**Supplementary Figure 2.**
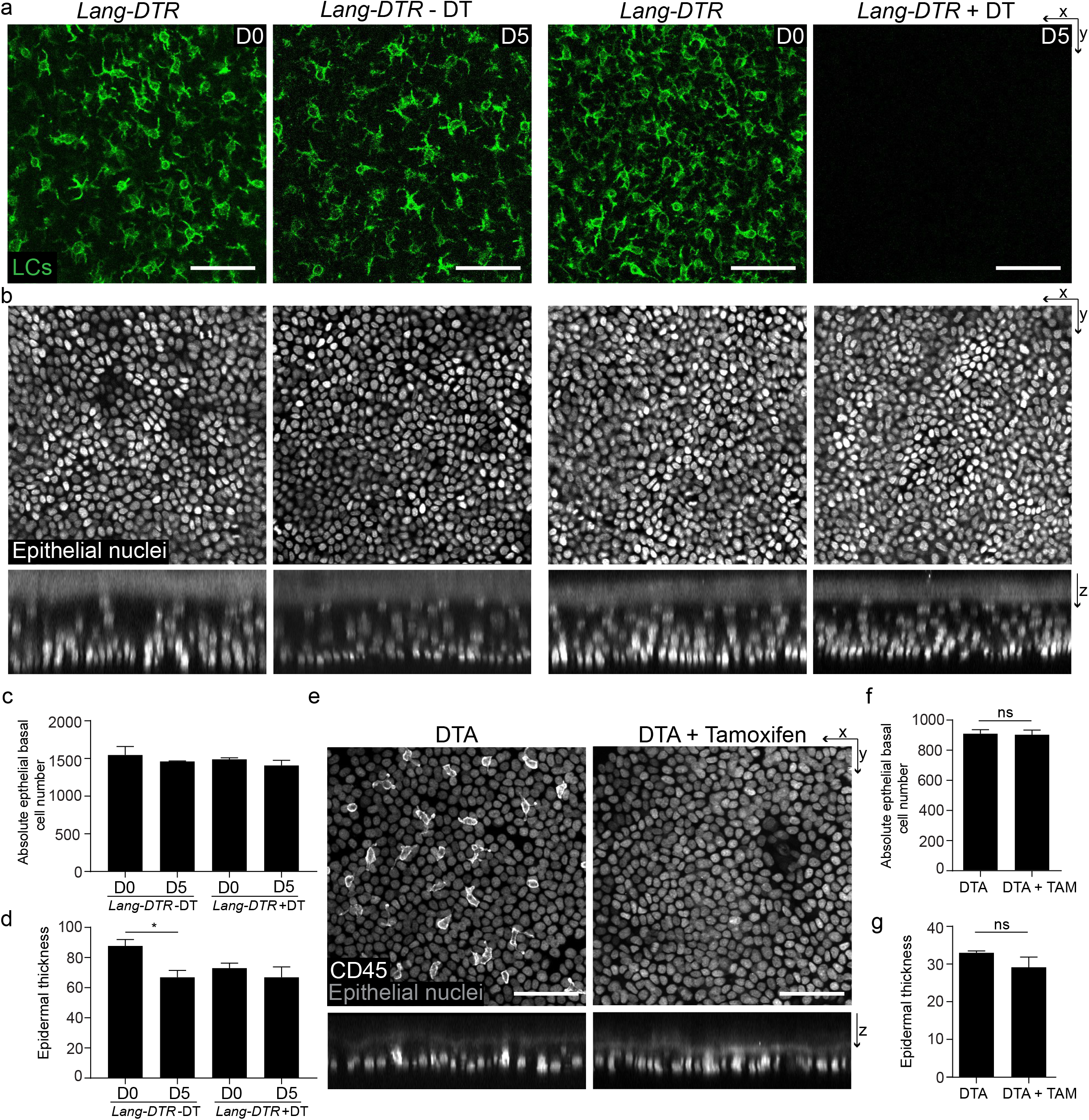
Full depletion of LCs and DETCs does not alter epidermal architecture. (a) X-y view of the basal layer of the epidermis in the paw with LCs in green (GFP) comparing *Lang-DTR* control mice without the addition of diphtheria toxin (or - DT) to *Lang-DTR* mice given 1 dose of 1 μg/body weight diphtheria toxin for the acute depletion of LCs 5 days before harvest (+ DT) (representative images from 2 mice each). (b) X-y view of the basal layer and x-z view of the epidermis with epithelial nuclei in white, comparing *Lang-DTR* mice either - DT or + DT (representative images from 3 mice each). The images show that the depletion of LCs does not impact the density or epidermal architecture of epithelial cells in the paw. Scale bar, 50 μm. (c) Quantifications of epithelial basal cell number comparing *Lang-DTR* mice - DT or + DT pre depletion (D0) and 5 days post depletion (D5). Unpaired Student’s *t*-test (n=2 mice respectively). (d) Quantification of epidermal thickness comparing *Lang-DTR* mice -DT or +DT at D0 and D5 with a small significance when comparing controls at D0 compared to D5 highlighting the variability in regions across mice. Area quantified 0.0625 mm^2^. \**p*<0.05, unpaired Student’s *t*-test (n=2 mice respectively). (e) X-y view of the basal layer and x-z view of the epidermis in the ear with epithelial nuclei in gray and immune cells (CD45+) in white comparing control *huLangerin-CreER; TCR δ-CreER; Rosa-GFP-stop-DTA* mice without tamoxifen (DTA) to *huLangerin-CreER; TCR δ-CreER; Rosa-GFP-stop-DTA* mice given 2 mgs of tamoxifen on 2 consecutive days 8 days before harvest (DTA + tamoxifen) (representative images from 2 mice each). The images show that the simultaneous depletion of both LCs and DETCs does not impact the density or epidermal architecture of epithelial cells in the ear. Scale bar, 50 μm. (f) Quantification of epithelial basal cell number comparing *huLangerin-CreER; TCR δ-CreER; Rosa-GFP-stop-DTA* either - Tamoxifen or + Tamoxifen at day 8. Unpaired Student’s *t*-test (n=2 mice respectively). (g) Quantification of epidermal thickness comparing *huLangerin-CreER; TCR δ-CreER; Rosa-GFP-stop-DTA* either - Tamoxifen or + Tamoxifen at day 8 showed that simultaneous depletion of LCs and DETCs did not significantly affect epidermal thickness. Area quantified 0.0625 mm^2^. Unpaired Student’s *t*-test (n=2 mice respectively).

**Supplementary Figure 3.**
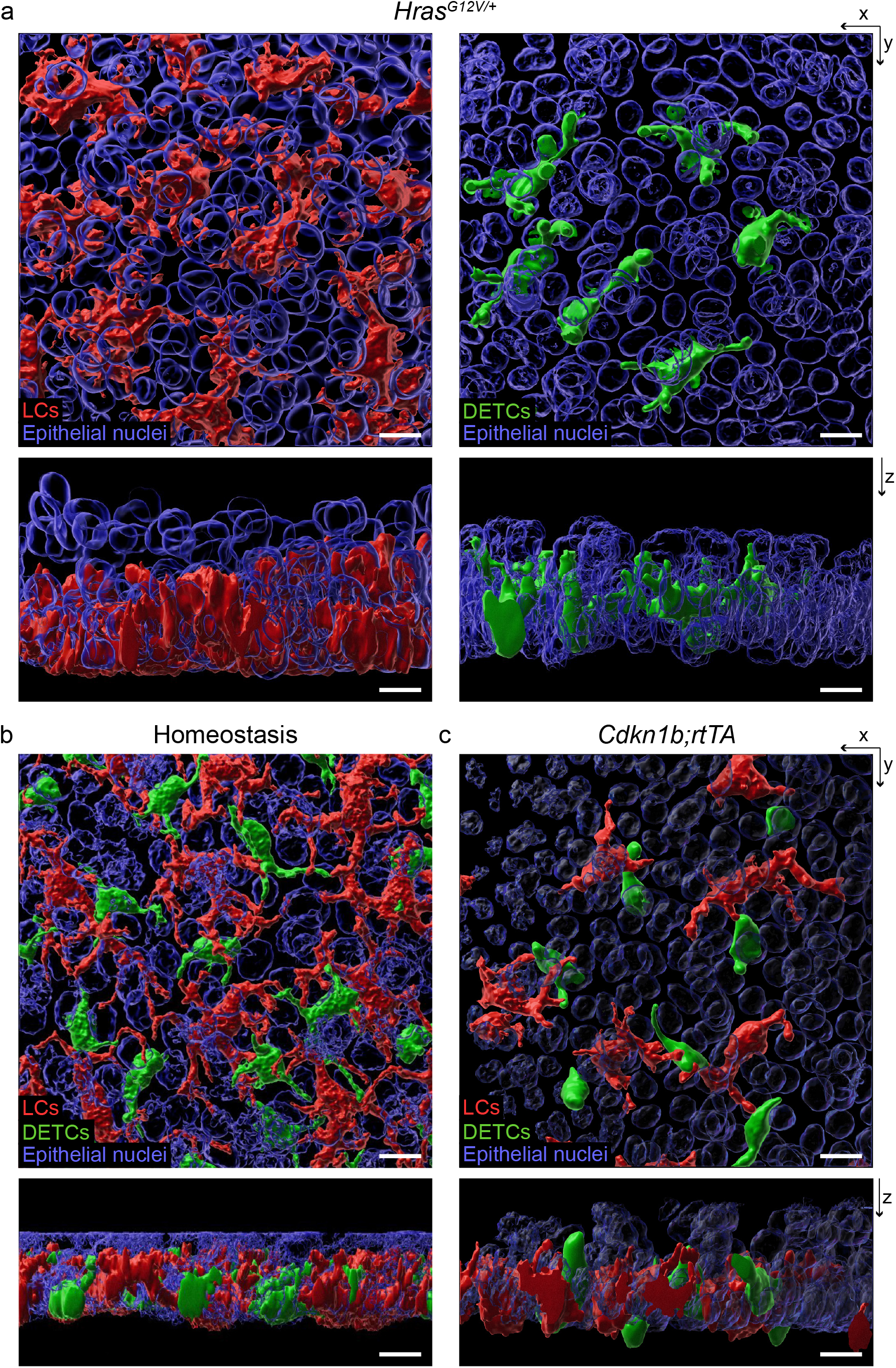
LCs and DETCs remain embedded in the basal layer of the ear epidermis upon changes in epithelial basal cell density (a-c) 3D reconstruction with Imaris surface rendering shows that LCs (red), DETCs (green), and epithelial cells (blue) are complicatedly intermixed in the epidermis. (a) X-y and x-z views show that LCs and DETCs embed in the basal layer of the epidermis in *Hras^G12V^* mice 6 weeks after 1 dose of 2 mgs tamoxifen similar to (b) WT LCs and DETCs in homeostasis and (c) *Cdkn1b;rtTA* mice 3 days after induction with 1 mg/ml doxycycline. Scale bar, 10 μm.

**Supplementary Figure 4.**
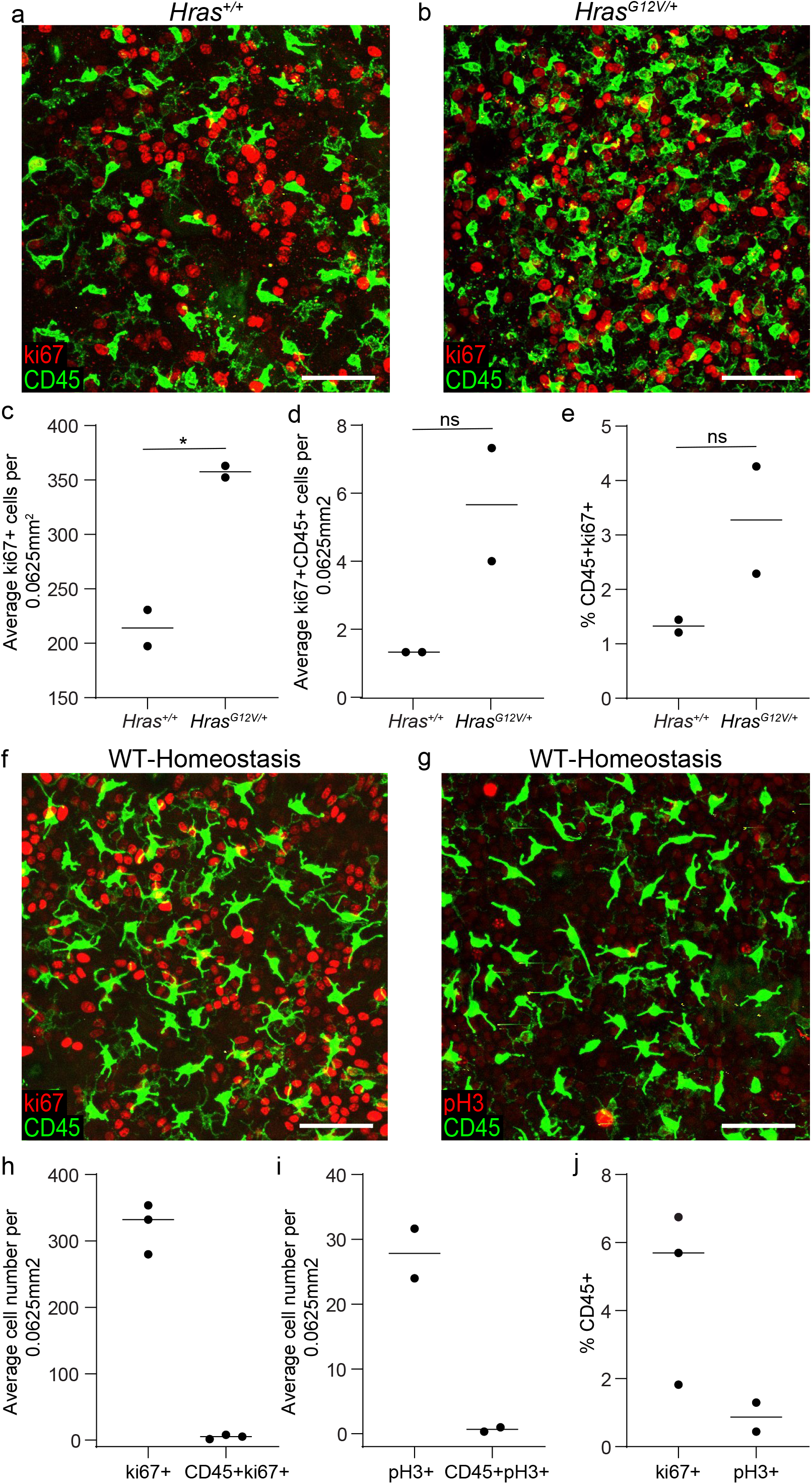
Proliferation in LCs and DETCs is enhanced in a model of increased epithelial and immune cell density. (a,b) Staining of the epidermal basal layer of the ear in *Hras^G12V/+^* compared to control *Hras ^+/+^* mice 6 weeks after induction with 1 dose of 2 mgs of tamoxifen for the proliferation marker ki67 (representative of 2 mice respectively). Scale bar, 50μm. (c-e) Quantifications revealed a 2-fold increase in the percentage of immune cells that are cycling (CD45+ki67+) in the *Hras^G12V/+^* mice compared to controls suggesting that immune cells in this model have enhanced proliferative capacity. Area quantified 0.0625 mm^2^ X 3 regions per mouse. Unpaired Student’s *t*-test (n=2 mice respectively). (f-j) Stainings of the epidermal basal layer of the ear in 6 week old CD1 mice for proliferation markers shown in red (f) ki67 and (g) phosphohistone H3 to enumerate proliferating immune cells (CD45+) shown in green (representative of 2 mice respectively). Scale bar, 50μm. (h-j) Quantifications show that an average of 5% of immune cells are cycling (CD45+ki67+) and 1% of immune cells are actively dividing (CD45+pH3+) during homeostasis. Area quantified 0.0625 mm^2^ X 3 regions per mouse. Unpaired Student’s *t*-test (n=3 mice for ki67 and n=2 mice for pH3)

**Supplementary Figure 5.**
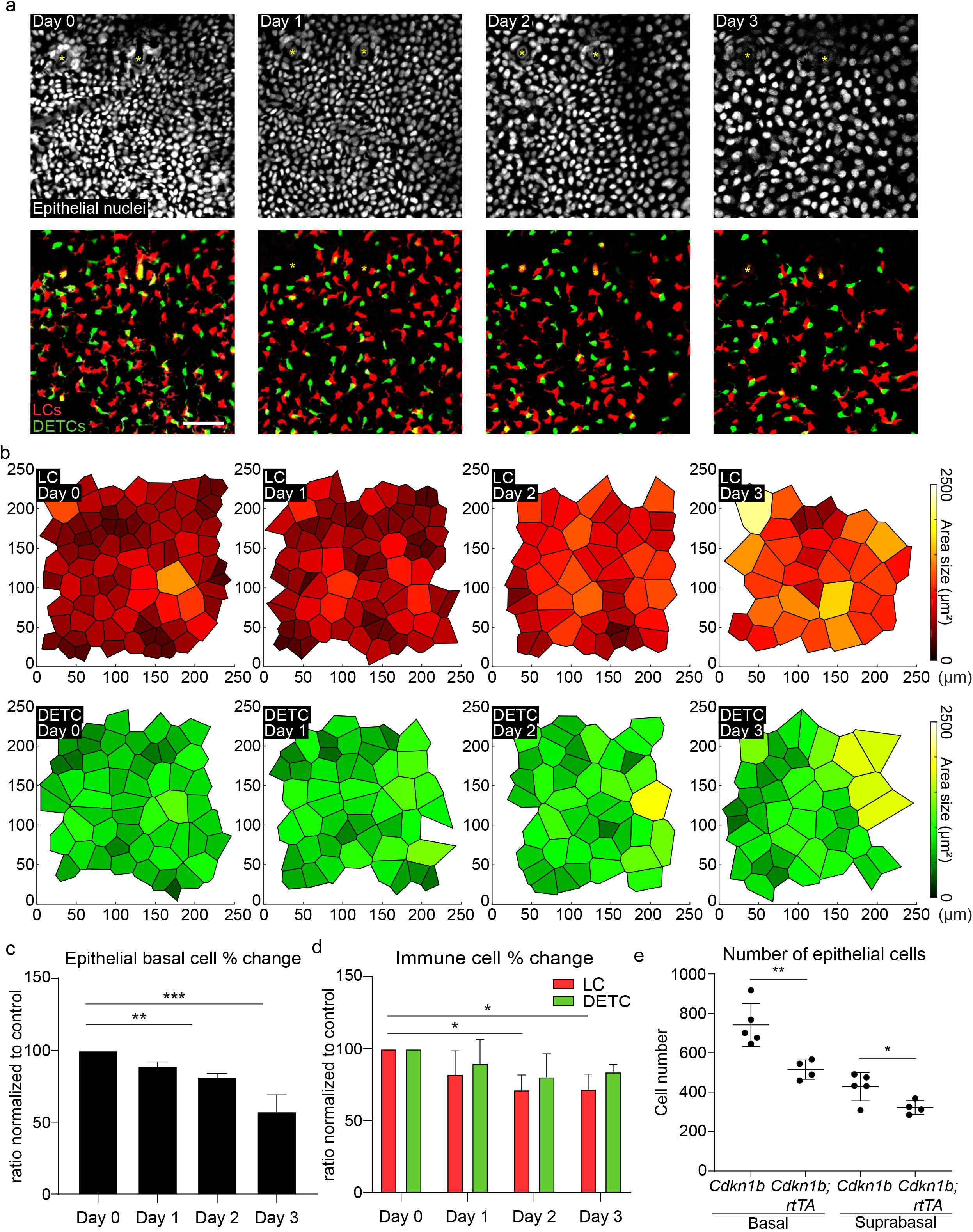
Immune cell density gradually changes along with changes in epithelial basal density. (a) The densities of LCs and DETCs in mice with blocked proliferation of epithelial stem cells *(Cdkn1b; K14-rtTA)* decrease in proportion to a decrease in the density of epithelial basal cells during a 3-day revisit experiment in as early as 1 day post induction with 1 mg/ml doxyclycline. Epithelial nuclei are white (top panel). LCs are red and DETCs are green (bottom panel). Yellow asterisk denotes hair follicle. (n=3 mice respectively). Scale bar, 50 μm. (b) Voronoi diagrams showing that the spatial distribution of LCs (top panel, red) and DETCs (bottom panel, green) is maintained during the 3 day revisit despite the decrease in cell number. (c) Quantification of epithelial basal cell number revealed a significant decrease over the timecourse. ** *p*<0.01 and ***p<0.001, unpaired Student’s t-test (n=3 mice respectively), (d) Quantification of LC and DETC number for both LCs and DETCs decreased over the timecourse. Area quantified 0.25 mm^2^. * *p*< 0.05, unpaired Student’s t-test (n=3 mice respectively). (e) Fiji software was used to count nucleated cells in the epidermis based on arbitrary distance from the SHG of the collagen in 260 × 260 ROIs of 40X images (0.27μm/pixel, 1μm step size). Basal cell counts comparing *Cdkn1b;rtTA* mice to *Cdkn1b* controls at day 3 post induction with 1 mg/ml doxycycline show a significant drop in epithelial basal density as expected. Suprabasal cell counts which include spinous and granular layers, comparing *Cdkn1b;rtTA* mice to *Cdkn1b* controls at day 3 post induction with 1 mg/ml doxycycline also show a significant drop in combined suprabasal cell density suggesting a coordination between the layers. Area quantified 0.0625 mm^2^. * *p*<0.05 and ** *p*<0.01, unpaired Student’s t-test (n=3 *Cdkn1b* mice and n=2 for *Cdkn1b;rtTA* mice).

**Supplementary Figure 6.**
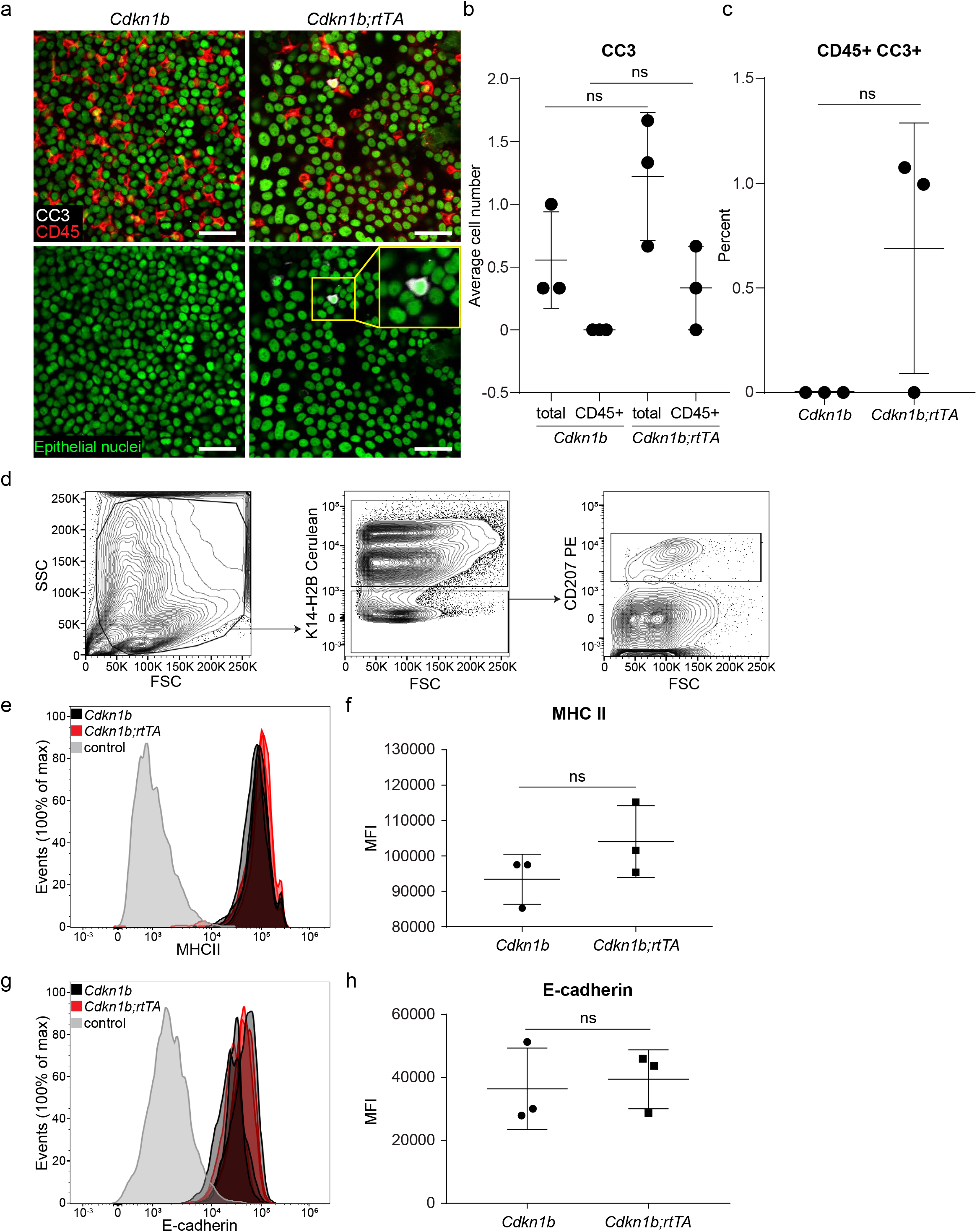
Cell death and activation do not contribute to the density drop of LCs and DETCs in *Cdkn1b;rtTA* mice. (a) Ears were taken from *Cdkn1b* and *Cdkn1b;rtTA* mice 3 days post induction with 1 mg/ml of doxycycline. Staining of the epithelial basal layer of the ear for the apoptotic marker cleaved-caspase 3 (CC3) shown in white of *Cdkn1b;rtTA* (right panels) and control *Cdkn1b* mice (left panels) (representative images of 3 mice respectively). LCs and DETCs are shown in red (CD45) and epithelial basal cell nuclei are shown in green. Inset in bottom right panel shows an epithelial cell positive for CC3. Scale bar, 40μm. (b,c) Quantifications of CC3 revealed very rare overall events and no significant difference in the percentage of immune cells undergoing apoptosis (CD45+CC3+) in *Cdkn1b;rtTA* compared to controls suggesting that cell death is not the cause of the decrease in LC and DETC density in this model. Area quantified 0.0625 mm^2^ X 3 regions per mouse. Unpaired Student’s t-test (n=3 mice respectively). (d) Epidermal single cell suspensions were processed from ear of mice 3 days post induction with 1mg/ml of doxycycline for flow cytometry and gated for LCs using anti-langerin antibody (CD207). K14-H2BCerulean was used to gate out the epithelial cells. (e,f) LCs from *Cdkn1b;rtTA* mice (closed red) show no significant difference in the expression levels of the known activation marker MHCII compared to those of *Cdkn1b* control mice (solid black) in the epidermis showing that LCs in *Cdkn1b;rtTA* mice are not activated. Negative control (gray) is gated on cells negative for the epithelial cell marker K14-H2BCerulean and immune cell markers for LCs (CD207) and DETCs (not shown). Unpaired Student’s t-test (n=3 mice respectively). (g-h) LCs from *Cdkn1b;rtTA* mice (closed red, n=3) also showed no significant difference compared to *Cdkn1b* control mice (closed black, n=3) in the expression levels of E-cadherin, a mediator of cell-cell interaction known to be down-regulated in activated LCs. Negative control (gray) is gated on cells negative for the epithelial cell marker K14-H2BCerulean and immune cell markers for LCs (CD207) and DETCs (not shown). Unpaired Student’s t-test (n= 3 mice respectively).

**Supplementary Figure 7.**
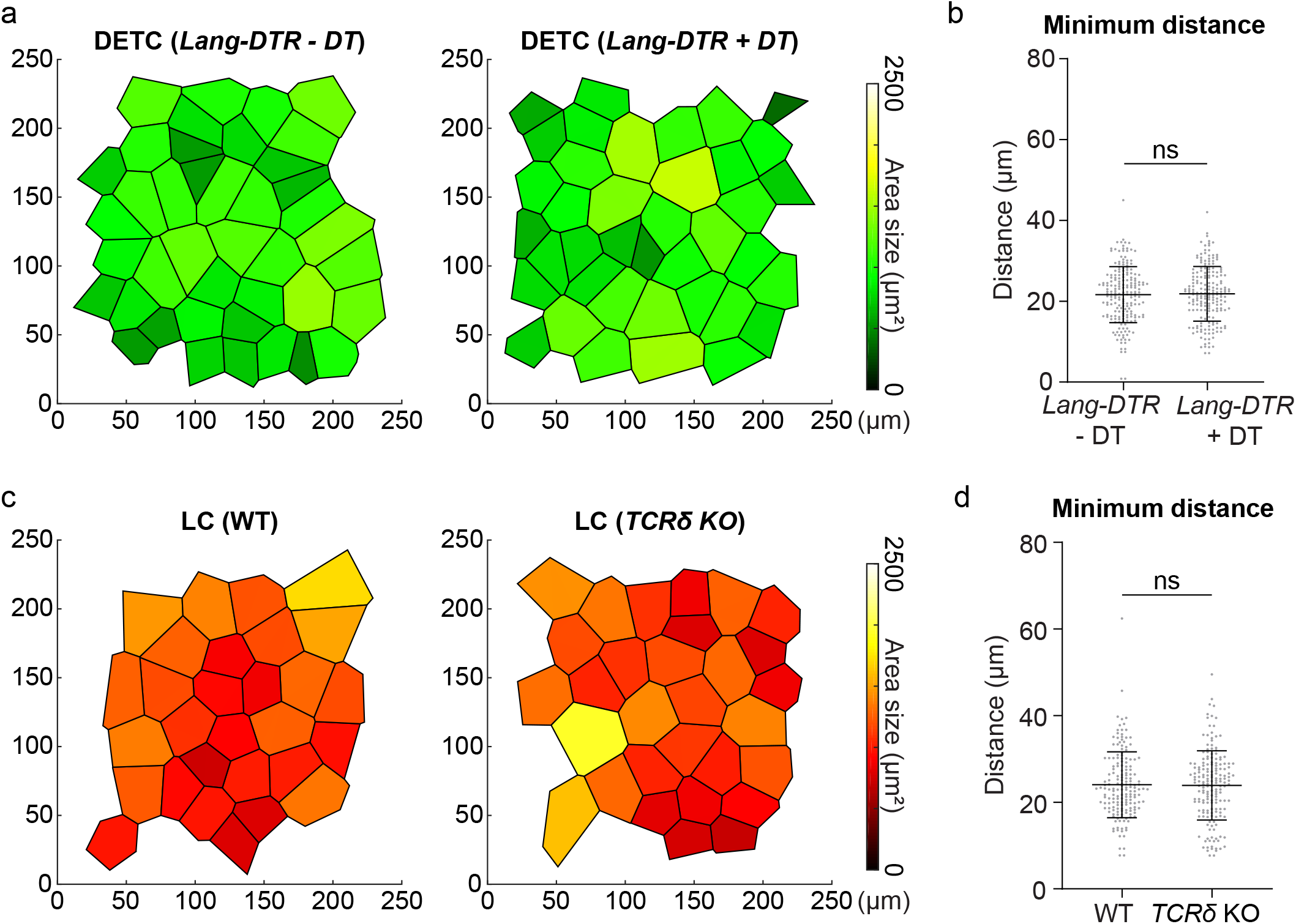
The distribution of one immune population is maintained upon the depletion of the other. (a,c) Voronoi diagrams were generated from images acquired for the quantifications in Supplementary Figure 1 comparing (a) the distribution of DETCs in *Lang-DTR* mice with and without one dose of 1 ug/body weight of diphtheria toxin 5 days post depletion or (c) that of LCs in *TCRδ KO* compared to WT controls showing that the distribution of the remaining immune population is not disrupted in either models. (b,d) Minimum distance quantifications showed no significant differences regardless of which immune population had been ablated when compared to controls showing that (b) DETC and (d) LC patterns are maintained in the absence of the other population. Area quantified 0.625 mm^2^. Unpaired Student’s t-test (n=3 mice respectively).

**Supplementary Figure 8.**
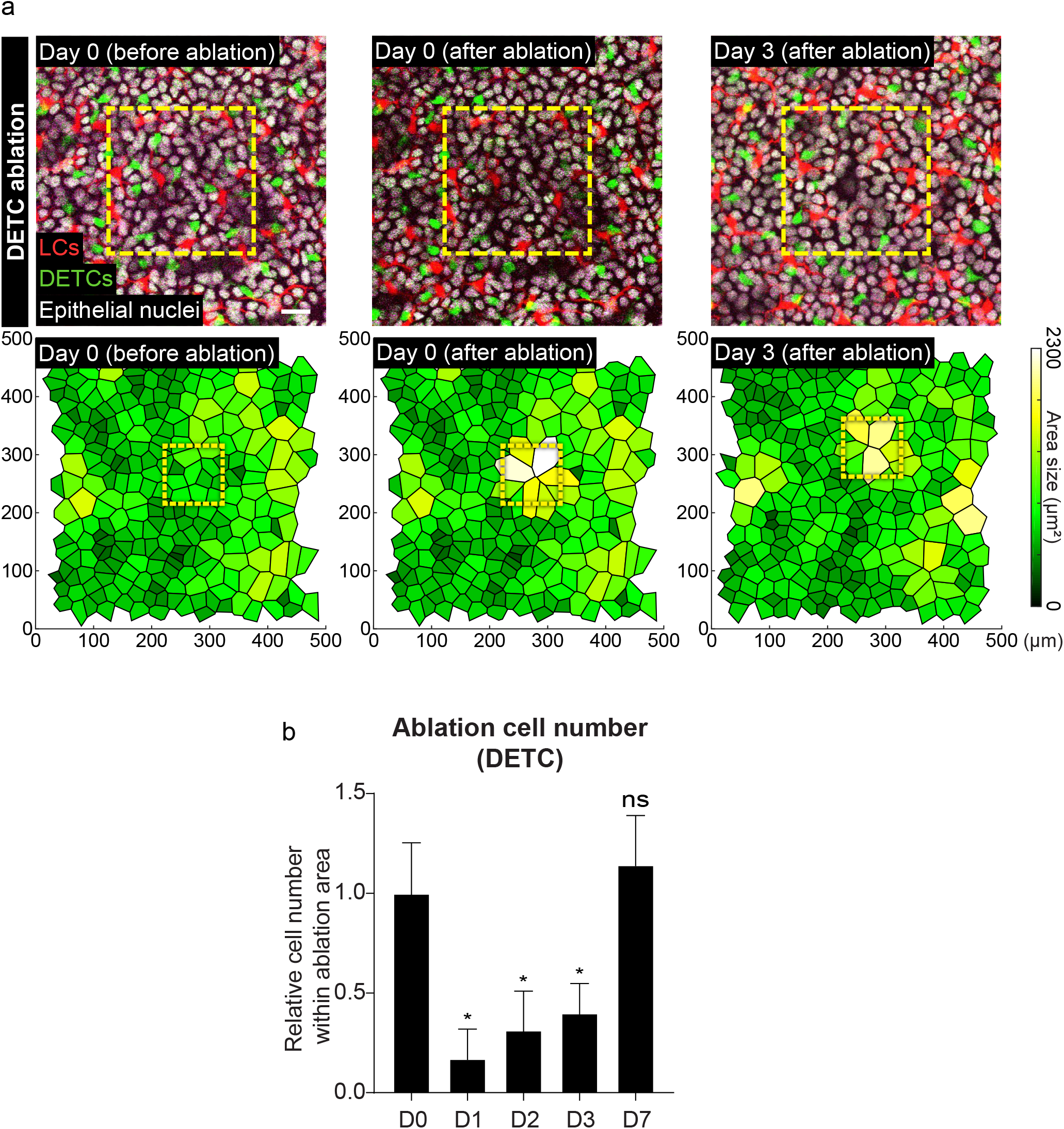
Surviving DETCs re-establish a regular pattern after local cell loss. (a) Local laser ablation of DETCs. DETCs within the yellow box (100 μm X 100 μm) are ablated by multiphoton laser (810 nm) and the same region is revisited 3 days after the ablation. The upper panel shows actual images from a mouse from day 0 before ablation to day 3 post ablation. Scale bar, 10μm. The lower panel displays the Voronoi diagram for DETCs generated from the images at each timepoint and encompasses a larger area around the ablation site. Neighboring DETCs migrate into the ablated region and re-establish a regular pattern. Area size 0.25 mm^2^ (representative images from 4 mice). (b) The DETC pattern within the ablated region was quantified from day 1 to day 7 and compared to the initial number at day 0. * *p*<0.05, unpaired Student’s *t*-test (n=4 mice respectively).

**Supplementary Figure 9.**
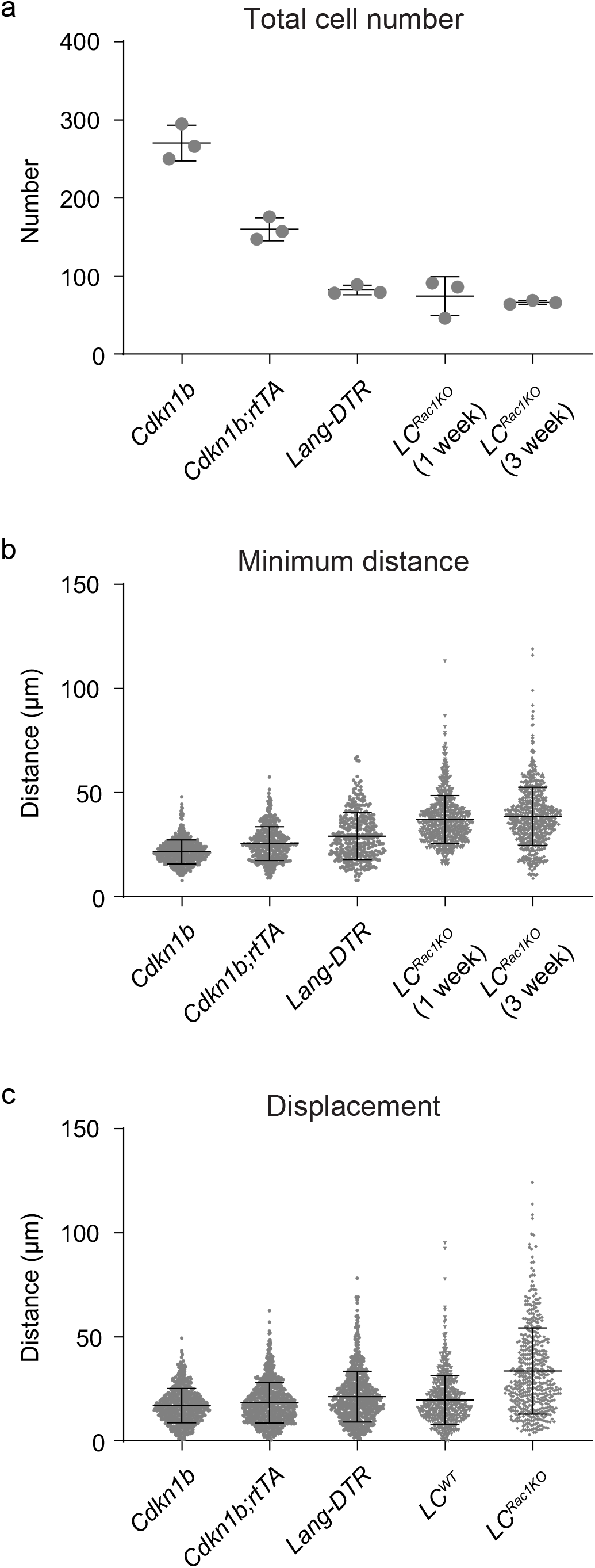
LCRac1^KO^ have increased mobility in the epidermis compared to models with similar LC density. (a-b) Total cell number and minimum distance was compared across models (a) Total cell number was quantified across the models to reveal similar densities of LCs in the Rac1 model (LC^Rac1KO^) compared to the LCs of *Cdkn1b;rtTA* and *Lang-DTR* mice (n=3 mice respectively). (b) The data show that LC^Rac1KO^ have increased variability in the minimum distance between neighbors compared to *Cdkn1b;rtTA* and *Lang-DTR* mice, suggesting that a drop in LC density is not sufficient to explain the disrupted pattern. Note, the data used for *Lang-DTR* is the same data set as in Figure 4d,e for Day 8 and the data used for LC^Rac1KO^ at 1 week and 3 weeks are the same data sets used in Figure 5f (n=3 mice respectively). (c) Displacement analysis shows a greater distance change in the relative position of LC^Rac1KO^ at day 0 versus day 3 compared to LCs of *Cdkn1b;rtTA* mice during 72 hours and LCs of *Lang-DTR* mice over the course of 8 days showing that LC^Rac1KO^ have less constrained mobility in the epidermis. The data used for LC^Rac1KO^ at 1 week and 3 weeks are the same data sets used in Figure 5b (n=3 mice respectively).

**Supplementary Figure 10.**
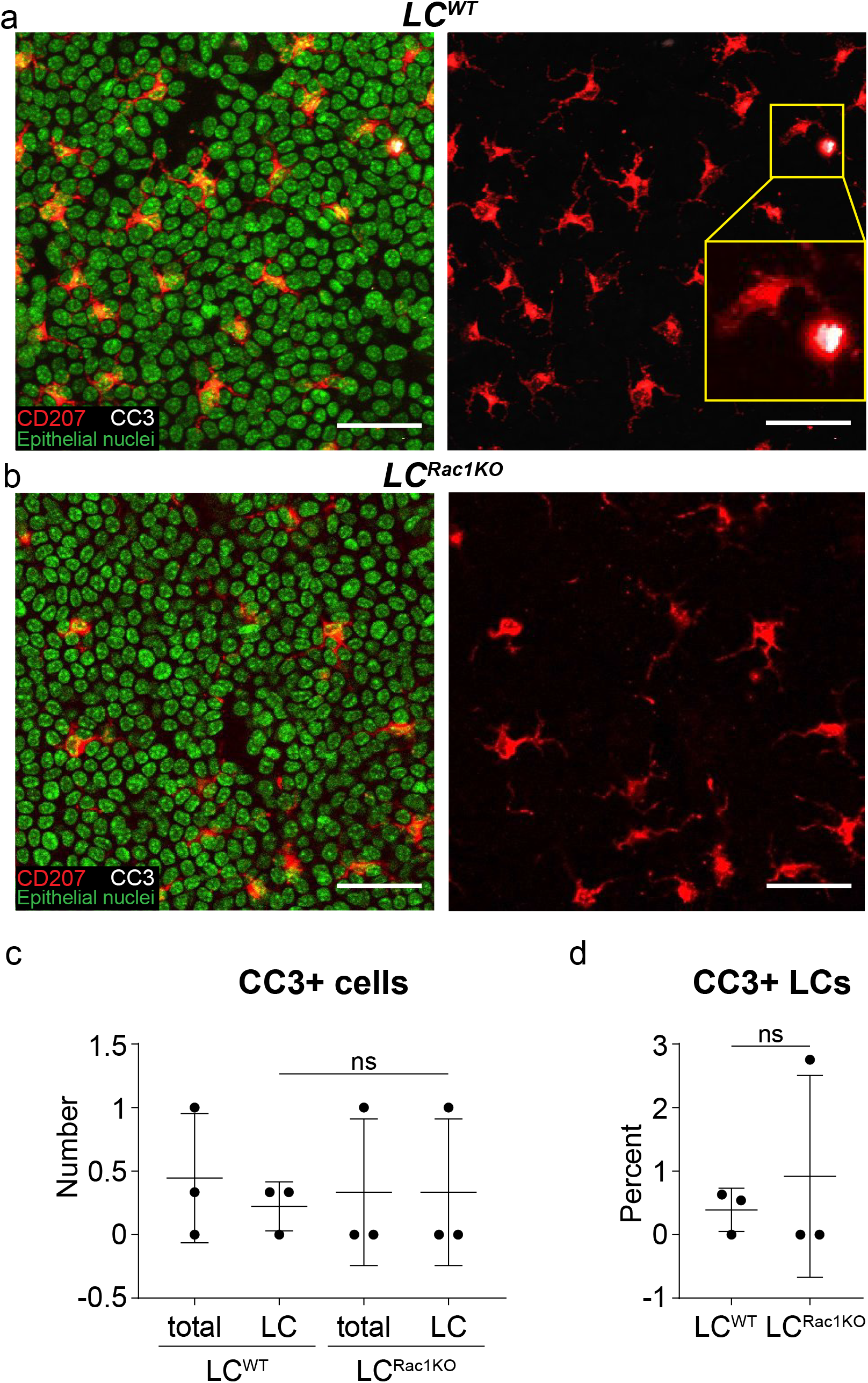
Cell death does not contribute to the density drop observed in the LC^Rac1KO^ phenotype. (a-b) Staining of the epithelial basal layer of the ear for the apoptotic marker cleaved-caspase (CC3) shown in white of (a) *huLangerin-CreER; Rac1^+/+^* mice and (b) *huLangerin-CreER; Rac1^fl/fl^* mice days post induction with 2 mg dose of tamoxifen given on 5 consecutive days. LCs are shown in red (CD207) and green (Hoechst) and epithelial basal cell nuclei are shown in green (Hoechst). Inset in bottom right panel shows an LC cell positive for CC3. (c,d) Quantifications show very few apoptotic events (< 1%) in the LCs of both *huLangerin-CreER; Rac1^+/+^* mice and *huLangerin-CreER; Rac1^+/+^* control mice (CC3+CD207+) showing that LCs are not dying in this model (data is representative n=3 for *huLangerin-CreER; Rac1^+/+^* mice, n=3 for *huLangerin-CreER; Rac1^fl/fl^* mice). Scale bar, 40 μm. Area quantified 0.0625 mm^2^ X 3 regions/mouse. Unpaired Student’s *t*-test (n=3 mice respectively).

**Supplementary Figure 11.**
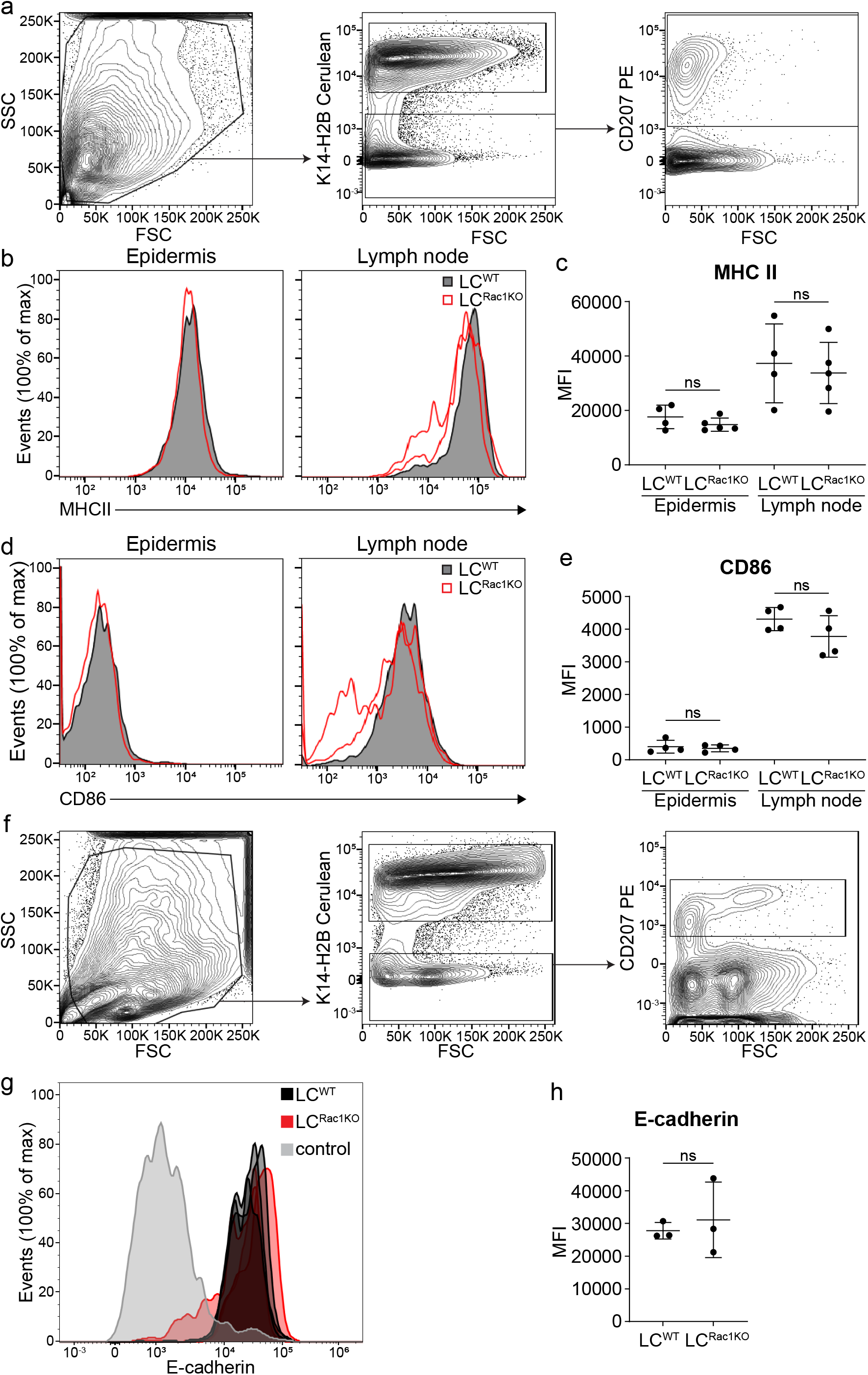
Activation does not contribute to the drop in density observed in LC^Rac1KO^. (a) Ears and lymph nodes were taken from mice 7 days post induction with 2mg dose of tamoxifen given on 5 consecutive days. Epidermal and lymph node single cell suspensions were processed for flow cytometry and gated for LCs using anti-langerin antibody (CD207). K14-H2BCerulean was used to gate out the epithelial cells. (b-e) LC^Rac1KO^ (open red) show no significant difference in the expression levels of the known activation markers MHCII and CD86 compared to LC^wt^(solid black) in either the epidermis or draining lymph nodes showing that LC^Rac1KO^ are not activated in this model. Unpaired Student’s *t*-test (data is representative n=4 for *huLangerin-CreER; Rac1^+/+^* mice, n=5 for *huLangerin-CreER; Rac1^fl/fl^* mice). (f) Epidermal cell suspensions were processed for flow cytometry and gated for LCs using anti-langerin antibody (CD207). K14-H2BCerulean was used to gate out the epithelial cells. (g-h) LC^Rac1KO^ (closed red, n=3) show no significant difference in the expression levels of the known mediator of cell-cell interaction E-cadherin compared to LC^wt^ (solid black, n=3). Negative control (gray) is gated on cells negative for the epithelial cell marker K14-H2BCerulean and immune cell markers for LCs (CD207) and DETCs (not shown). Unpaired Student’s *t*-test (data is representative n=3 for *huLangerin-CreER; Rac1^+/+^* mice, n=3 for *huLangerin-CreER; Rac1^fl/fl^* mice).

## Supplementary Video Legends

**Supplementary Video 1. 3D reconstruction of epidermis.** 3D reconstruction with Imaris surface rendering shows that LCs (red), DETCs (green), and epithelial cells (white) are complicatedly intermixed in the epidermis. LCs and DETCs extrude their dendrite vertically to the surface of the epidermis as well as laterally between the epithelial cells. Scale bar, 40 μm.

**Supplementary Video 2. Dendritic movement of LCs and DETCs during homeostasis.**Time-lapse recording over 4 hours of LCs and DETCs shows stable cell bodies while their dendrites dynamically explore junctional spaces between neighboring epithelial basal cells. LCs (red), DETCs (green) and nuclei of epithelial stem cells (white). Yellow arrows point at dendritic activity in the junctional space of epithelial basal cells. Scale bar, 50 μm.

**Supplementary Video 3. Time-lapse movies of LC^WT^ and LC^Rac1KO^.**Time-lapse recording over 4 hours of LC^WT^ and LC^Rac1KO^ shows that LCs are stable in position and protrude dendrites over time regardless of the Rac1 KO in the LCs. Yellow arrows point at dendritic activity. Zoom in movies repeat 5 times. Scale bar of wide view, 50 μm. Scale bar of zoom in, 20 μm.

## Supplementary Table Legends

**Supplementary Table 1.**Statistics source data for Figure 1e, 1f, 1g, 2b, 2c, 2d, 2f, 2g, 2h, 2j, 2k, 2l, 3b, 3c, 3d, 3e, 3f, 3g, 4a, 4b, 4c, 4d, 4e, 4f, 4g, 5b, 5d, 5e, 5f, and for Supplementary Figure 1c, 1d, 1e, 1f, 1g, 1h, 2c, 2d, 2f, 2g, 4c, 4d, 4e, 4h, 4i, 4j, 5b, 5c, 5d, 5e, 6b, 6c, 6f, 6h, 7a, 7b, 7c, 7d, 8a, 8b, 9a, 9b, 9c, 10c, 10d, 11c, 11e and 11h.

## Rebuttal Video Legends

**Rebuttal Video 1. LCs re-populate in the absence of LC divisions in a partial depletion.**Timelapse recordings of a partial depletion (2 ng/g of body weight) of LCs in *Lang-DTR* mice over 3 hours captured real time dynamics of LCs during repatterning 7 days after depletion and illustrate their outward movement towards regions devoid of LCs in the absence of LC divisions. LCs (red) and nuclei of epithelial basal cells (green). Area size 0.75 mm^2^.

**Rebuttal Video 2. LCs re-populate in the absence of LC divisions in a full depletion.**Timelapse recordings of a full depletion (1 μg/ body weight) of LCs in *Lang-DTR* mice over 4 hours captured real time dynamics of LCs during repatterning 12 days after depletion and illustrate their outward movement towards regions devoid of LCs in the absence of LC divisions. LCs (red) and nuclei of epithelial basal cells (green). Area size 1mm^2^.

## Notes

### Competing Interest Statement

The authors have declared no competing interest.

## References

1. Gonzales, K.A.U. and E. Fuchs, Skin and Its Regenerative Powers: An Alliance between Stem Cells and Their Niche. Dev Cell, 2017. 43(4): p. 387–401.

2. Park, S., V. Greco, and K. Cockburn, Live imaging of stem cells: answering old questions and raising new ones. Curr Opin Cell Biol, 2016. 43: p. 30–37.

3. Simpson, C.L., D.M. Patel, and K.J. Green, Deconstructing the skin: cytoarchitectural determinants of epidermal morphogenesis. Nat Rev Mol Cell Biol, 2011. 12(9): p. 565–80.

4. Solanas, G. and S.A. Benitah, Regenerating the skin: a task for the heterogeneous stem cell pool and surrounding niche. Nat Rev Mol Cell Biol, 2013. 14(11): p. 737–48.

5. Tai, K., K. Cockburn, and V. Greco, Flexibility sustains epithelial tissue homeostasis. Curr Opin Cell Biol, 2019. 60: p. 84–91.

6. Xin, T., V. Greco, and P. Myung, Hardwiring Stem Cell Communication through Tissue Structure. Cell, 2016. 164(6): p. 1212–1225.

7. Ho, A.W. and T.S. Kupper, T cells and the skin: from protective immunity to inflammatory skin disorders. Nat Rev Immunol, 2019. 19(8): p. 490–502.

8. Pasparakis, M., I. Haase, and F.O. Nestle, Mechanisms regulating skin immunity and inflammation. Nat Rev Immunol, 2014. 14(5): p. 289–301.

9. Tay, S.S., et al., The Skin-Resident Immune Network. Curr Dermatol Rep, 2014. 3: p. 13–22.

10. Brand, A., et al., E-Cadherin is Dispensable to Maintain Langerhans Cells in the Epidermis. J Invest Dermatol, 2020. 140(1): p. 132–142 e3.

11. Dainichi, T., et al., The epithelial immune microenvironment (EIME) in atopic dermatitis and psoriasis. Nat Immunol, 2018. 19(12): p. 1286–1298.

12. Eming, S.A., T. Krieg, and J.M. Davidson, Inflammation in wound repair: molecular and cellular mechanisms. J Invest Dermatol, 2007. 127(3): p. 514–25.

13. Jameson, J., et al., A role for skin gammadelta T cells in wound repair. Science, 2002. 296(5568): p. 747–9.

14. Kaplan, D.H., Ontogeny and function of murine epidermal Langerhans cells. Nat Immunol, 2017. 18(10): p. 1068–1075.

15. Leoni, G., et al., Wound repair: role of immune-epithelial interactions. Mucosal Immunol, 2015. 8(5): p. 959–68.

16. Lin, X., et al., An Ectoderm-Derived Myeloid-like Cell Population Functions as Antigen Transporters for Langerhans Cells in Zebrafish Epidermis. Dev Cell, 2019. 49(4): p. 605–617 e5.

17. Merad, M., F. Ginhoux, and M. Collin, Origin, homeostasis and function of Langerhans cells and other langerin-expressing dendritic cells. Nat Rev Immunol, 2008. 8(12): p. 935–47.

18. Nielsen, M.M., D.A. Witherden, and W.L. Havran, gammadelta T cells in homeostasis and host defence of epithelial barrier tissues. Nat Rev Immunol, 2017. 17(12): p. 733–745.

19. Sere, K., et al., Two distinct types of Langerhans cells populate the skin during steady state and inflammation. Immunity, 2012. 37(5): p. 905–16.

20. Strbo, N., N. Yin, and O. Stojadinovic, Innate and Adaptive Immune Responses in Wound Epithelialization. Adv Wound Care (New Rochelle), 2014. 3(7): p. 492–501.

21. Takashima, A. and P.R. Bergstresser, Cytokine-mediated communication by keratinocytes and Langerhans cells with dendritic epidermal T cells. Semin Immunol, 1996. 8(6): p. 333–9.

22. Lewis, J.M., et al., Mechanisms of chemical cooperative carcinogenesis by epidermal Langerhans cells. J Invest Dermatol, 2015. 135(5): p. 1405–1414.

23. Lewis, J.M., et al., Langerhans Cells Facilitate UVB-Induced Epidermal Carcinogenesis. J Invest Dermatol, 2015. 135(11): p. 2824–2833.

24. Naik, S., et al., Inflammatory memory sensitizes skin epithelial stem cells to tissue damage. Nature, 2017. 550(7677): p. 475–480.

25. Sugita, K., et al., Innate immunity mediated by epidermal keratinocytes promotes acquired immunity involving Langerhans cells and T cells in the skin. Clin Exp Immunol, 2007. 147(1): p. 176–83.

26. Bernard, J.J., R.L. Gallo, and J. Krutmann, Photoimmunology: how ultraviolet radiation affects the immune system. Nat Rev Immunol, 2019. 19(11): p. 688–701.

27. Nakatsuji, T., et al., Staphylococcus aureus Exploits Epidermal Barrier Defects in Atopic Dermatitis to Trigger Cytokine Expression. J Invest Dermatol, 2016. 136(11): p. 2192–2200.

28. Chodaczek, G., et al., Body-barrier surveillance by epidermal gammadelta TCRs. Nat Immunol, 2012. 13(3): p. 272–82.

29. Kitashima, D.Y., et al., Langerhans Cells Prevent Autoimmunity via Expansion of Keratinocyte Antigen-Specific Regulatory T Cells. EBioMedicine, 2018. 27: p. 293–303.

30. Kubo, A., K. Nagao, and M. Amagai, 3D visualization of epidermal Langerhans cells. Methods Mol Biol, 2013. 961: p. 119–27.

31. Kubo, A., et al., External antigen uptake by Langerhans cells with reorganization of epidermal tight junction barriers. J Exp Med, 2009. 206(13): p. 2937–46.

32. Matsui, T. and M. Amagai, Dissecting the formation, structure and barrier function of the stratum corneum. Int Immunol, 2015. 27(6): p. 269–80.

33. Nagao, K., et al., Stress-induced production of chemokines by hair follicles regulates the trafficking of dendritic cells in skin. Nat Immunol, 2012. 13(8): p. 744–52.

34. Yoshida, K., et al., Distinct behavior of human Langerhans cells and inflammatory dendritic epidermal cells at tight junctions in patients with atopic dermatitis. J Allergy Clin Immunol, 2014. 134(4): p. 856–64.

35. Bobr, A., et al., Autocrine/paracrine TGF-beta1 inhibits Langerhans cell migration. Proc Natl Acad Sci U S A, 2012. 109(26): p. 10492–7.

36. Madisen, L., et al., A robust and high-throughput Cre reporting and characterization system for the whole mouse brain. Nat Neurosci, 2010. 13(1): p. 133–40.

37. Jung, S., et al., Analysis of fractalkine receptor CX(3)CR1 function by targeted deletion and green fluorescent protein reporter gene insertion. Mol Cell Biol, 2000. 20(11): p. 4106–14.

38. Mesa, K.R., et al., Homeostatic Epidermal Stem Cell Self-Renewal Is Driven by Local Differentiation. Cell Stem Cell, 2018. 23(5): p. 677–686 e4.

39. Xin, T., et al., Flexible fate determination ensures robust differentiation in the hair follicle. Nat Cell Biol, 2018. 20(12): p. 1361–1369.

40. Rompolas, P., et al., Live imaging of stem cell and progeny behaviour in physiological hairfollicle regeneration. Nature, 2012. 487(7408): p. 496–9.

41. Rompolas, P., et al., Spatiotemporal coordination of stem cell commitment during epidermal homeostasis. Science, 2016. 352(6292): p. 1471–4.

42. Pineda, C.M., et al., Intravital imaging of hair follicle regeneration in the mouse. Nat Protoc, 2015. 10(7): p. 1116–30.

43. Rompolas, P., K.R. Mesa, and V. Greco, Spatial organization within a niche as a determinant of stem-cell fate. Nature, 2013. 502(7472): p. 513–8.

44. Kissenpfennig, A., et al., Dynamics and function of Langerhans cells in vivo: dermal dendritic cells colonize lymph node areas distinct from slower migrating Langerhans cells. Immunity, 2005. 22(5): p. 643–54.

45. Itohara, S., et al., T cell receptor delta gene mutant mice: independent generation of alpha beta T cells and programmed rearrangements of gamma delta TCR genes. Cell, 1993. 72(3): p. 337–48.

46. Bobr, A., et al., Acute ablation of Langerhans cells enhances skin immune responses. J Immunol, 2010. 185(8): p. 4724–8.

47. De Creus, A., et al., Langerhans cells that have matured in vivo in the absence of T cells are fully capable of inducing a helper CD4 as well as a cytotoxic CD8 response. J Immunol, 2000. 165(2): p. 645–53.

48. Taveirne, S., et al., Langerhans cells are not required for epidermal Vgamma3 T cell homeostasis and function. J Leukoc Biol, 2011. 90(1): p. 61–8.

49. Zhang, B., et al., Differential Requirements of TCR Signaling in Homeostatic Maintenance and Function of Dendritic Epidermal T Cells. J Immunol, 2015. 195(9): p. 4282–91.

50. Ivanova, A., et al., In vivo genetic ablation by Cre-mediated expression of diphtheria toxin fragment A. Genesis, 2005. 43(3): p. 129–35.

51. Brown, S., et al., Correction of aberrant growth preserves tissue homeostasis. Nature, 2017. 548(7667): p. 334–337.

52. Pineda, C.M., et al., Hair follicle regeneration suppresses Ras-driven oncogenic growth. J Cell Biol, 2019. 218(10): p. 3212–3222.

53. Pruitt, S.C., et al., Cdkn1b overexpression in adult mice alters the balance between genome and tissue ageing. Nat Commun, 2013. 4: p. 2626.

54. Xie, W., et al., Conditional expression of the ErbB2 oncogene elicits reversible hyperplasia in stratified epithelia and up-regulation of TGFalpha expression in transgenic mice. Oncogene, 1999. 18(24): p. 3593–607.

55. Park, S., et al., Tissue-scale coordination of cellular behaviour promotes epidermal wound repair in live mice. Nat Cell Biol, 2017. 19(2): p. 155–163.

56. Bauer, J., et al., A strikingly constant ratio exists between Langerhans cells and other epidermal cells in human skin. A stereologic study using the optical disector method and the confocal laser scanning microscope. J Invest Dermatol, 2001. 116(2): p. 313–8.

57. Numahara, T., et al., Spatial data analysis by epidermal Langerhans cells reveals an elegant system. J Dermatol Sci, 2001. 25(3): p. 219–28.

58. Sandrock, I., et al., Genetic models reveal origin, persistence and non-redundantfunctions of IL-17-producing gammadelta T cells. J Exp Med, 2018. 215(12): p. 3006–3018.

59. Gentek, R., et al., Epidermal gammadelta T cells originate from yolk sac hematopoiesis and clonally self-renew in the adult. J Exp Med, 2018. 215(12): p. 2994–3005.

60. Ghigo, C., et al., Multicolor fate mapping of Langerhans cell homeostasis. J Exp Med, 2013. 210(9): p. 1657–64.

61. Merad, M., et al., Langerhans cells renew in the skin throughout life under steady-state conditions. Nat Immunol, 2002. 3(12): p. 1135–41.

62. Marsh, E., et al., Positional Stability and Membrane Occupancy Define Skin Fibroblast Homeostasis In Vivo. Cell, 2018. 175(6): p. 1620–1633 e13.

63. Swetman, C.A., et al., Extension, retraction and contraction in the formation of a dendritic cell dendrite: distinct roles for Rho GTPases. Eur J Immunol, 2002. 32(7): p. 2074–83.

64. Grueber, W.B. and A. Sagasti, Self-avoidance and tiling: Mechanisms of dendrite and axon spacing. Cold Spring Harb Perspect Biol, 2010. 2(9): p. a001750.

65. Zipursky, S.L. and W.B. Grueber, The molecular basis of self-avoidance. Annu Rev Neurosci, 2013. 36: p.547–68.

66. Glogauer, M., et al., Rac1 deletion in mouse neutrophils has selective effects on neutrophil functions. J Immunol, 2003. 170(11): p. 5652–7.

67. Hotulainen, P. and C.C. Hoogenraad, Actin in dendritic spines: connecting dynamics to function. J Cell Biol, 2010. 189(4): p. 619–29.

68. Dogterom, M. and G.H. Koenderink, Actin-microtubule crosstalk in cell biology. Nat Rev Mol Cell Biol, 2019. 20(1): p. 38–54.

69. Luckashenak, N., et al., Rho-family GTPase Cdc42 controls migration of Langerhans cells in vivo. J Immunol, 2013. 190(1): p. 27–35.

70. Nishibu, A., et al., Behavioral responses of epidermal Langerhans cells in situ to local pathological stimuli. J Invest Dermatol, 2006. 126(4): p. 787–96.

71. Mohammed, J., et al., Stromal cells control the epithelial residence of DCs and memory T cells by regulated activation of TGF-beta. Nat Immunol, 2016. 17(4): p. 414–21.

72. Van den Bossche, J., et al., Regulation and function of the E-cadherin/catenin complex in cells of the monocyte-macrophage lineage and DCs. Blood, 2012. 119(7): p. 1623–33.

73. Mayumi, N., et al., E-cadherin interactions are required for Langerhans cell differentiation. Eur J Immunol, 2013. 43(1): p. 270–80.

74. Vasioukhin, V., et al., The magical touch: genome targeting in epidermal stem cells induced by tamoxifen application to mouse skin. Proc Natl Acad Sci U S A, 1999. 96(15): p. 8551–6.

75. Peron, S.P., et al., A Cellular Resolution Map of Barrel Cortex Activity during Tactile Behavior. Neuron, 2015. 86(3): p. 783–99.

76. Chen, X., et al., Endogenous expression of Hras(G12V) induces developmental defects and neoplasms with copy number imbalances of the oncogene. Proc Natl Acad Sci U S A, 2009. 106(19): p. 7979–84.

77. Jung, S., et al., In vivo depletion of CD11c+ dendritic cells abrogates priming of CD8+ T cells by exogenous cell-associated antigens. Immunity, 2002. 17(2): p. 211–20.

